# Pulsed stimulated Brillouin microscopy enables high-sensitivity mechanical imaging of live and fragile biological specimens

**DOI:** 10.1101/2022.11.10.515835

**Authors:** Fan Yang, Carlo Bevilacqua, Sebastian Hambura, Ana Neves, Anusha Gopalan, Koki Watanabe, Matt Govendir, Maria Bernabeu, Jan Ellenberg, Alba Diz-Muñoz, Simone Köhler, Georgia Rapti, Martin Jechlinger, Robert Prevedel

## Abstract

Brillouin microscopy is an emerging optical elastography technique capable of assessing mechanical properties of biological samples in a 3D, all-optical and hence non-contact fashion. The typically weak Brillouin scattering signal can be substantially enhanced via a stimulated photon-phonon process, which improves the signal-to-background ratio (SBR) as well as provides higher spectral resolution. However, current implementations of stimulated Brillouin spectroscopy (SBS) require high pump powers, which prohibit applications in many areas of biology, especially when studying photosensitive samples, or when live-imaging in 3D and/or over extended time periods. Here, we present a pulsed SBS scheme that takes full advantage of the non-linearity of the pump-probe interaction in SBS. In particular, we show that through quasi-pulsing and diligent optimization of signal detection parameters, the required pump laser power can be decreased ~20-fold without affecting the signal levels or spectral precision. Moreover, we devise a custom analysis approach that facilitates the analysis of complex, multi-peaked Brillouin spectra in order to harness the high spectral resolution of SBS for the specific identification of biomechanical components inside the point-spread function of the microscope. We then demonstrate the low-phototoxicity and high-specificity of our pulsed SBS approach by imaging sensitive single cells, zebrafish larvae, and mouse embryos as well as adult *C. elegans* with sub-cellular detail. Furthermore, our method permits observing the mechanics of organoids and *C. elegans* embryos over time. We expect that the substantially lower photo-burden and improved SBR of pulsed SBS will facilitate studying biomechanics in 3D at high spatio-temporal resolution in living biological specimens in a non-invasive manner, opening up exciting new possibilities for the field of mechanobiology.

---

At every length scale, the mechanical properties of tissues and cells play an intricate role in determining biological function. On the cellular scale, mechanical properties determine how cells respond to physical forces and their environment. On larger length scales, mechanical properties of tissues are thought to be imperative in the onset and progression of many diseases, such as eye disease, cancer and atherosclerosis^1–3^. Over the past decades, tremendous progress has been made in the fields of biomechanics to understand the relationship between mechanical forces, material properties and biochemical signals to regulate cellular function and tissue organization. However, while molecular and genetic components can be visualized through various fluorescence based imaging tools, assessing the mechanical, i.e. elastic and viscous, properties of living cells with similar spatio-temporal resolution has long been an open challenge. Existing biophysical techniques^3–5^ currently used in the field exhibit intrinsic limitations in this respect, as they are predominantly based on direct physical contact and involve direct application of external forces to the sample. Therefore, they are either limited to sample surfaces or lack the required 3D cellular resolution^3^.

Over the past decade, a new type of optical elastography, namely Brillouin microscopy (BM), has emerged as a non-destructive, label- and contact-free technique that allows probing of mechanical properties in a non-contact, label-free and high-resolution fashion^6,7^. It is based on light scattering of visible or infrared monochromatic (laser) light from gigahertz-frequency longitudinal acoustic phonons that are characteristic of the mechanical components of the material. The resulting Brillouin spectrum, i.e. the frequency shift (Ω_*B*_), and linewidth (*Γ_B_*) of the inelastically scattered light then provides information on the speed and attenuation of hypersonic acoustic waves in a (bio-)material from which visco-elastic properties can be derived. When coupled to a confocal microscope^6^, BM can achieve diffraction-limited resolution in 3D which has enabled a wide range of applications in biology, including studies of intracellular biomechanics in living cells^8^, of ex-vivo^9,10^ and in-vivo tissues^11–13^ and their components^14^ (such as collagen, elastin) as well as biomaterials^15^. As such it promises applications in early diagnosis of diseases (such as cancer, keratoconus)^16,17^. The typical signal levels in spontaneous Brillouin scattering, however, are relatively weak, due to the low scattering cross section of this process (~10^-10^ – 10^-12^), which has hampered the development of high-speed and/or high-sensitivity Brillouin microscopes.

Recently, stimulated Brillouin scattering (SBS) has been introduced and shown to enable high mechanical specificity as well as practical acquisition times of biological samples in Brillouin microscopy of biological samples^18,19^. The technique is based on coherently driving, and thus enhancing, the phonon population inside a sample at a given frequency. This can be achieved by creating an interference fringe pattern inside the sample via two overlapping laser (pump) beams^20^. This in turn ‘imprints’ an acoustic wave through electrostriction. By spatially overlapping and scanning a frequency-tunable ‘probe’ laser beam over the Brillouin spectrum the probe intensity (I_2_) at frequency ω_2_ experiences a stimulated Brillouin gain (ΔI_2_, SBG) or loss (SBL). In contrast to spontaneous Brillouin scattering, SBS enables spectral measurements at a higher resolution that are free of elastic background contributions, and thus enabling high signal levels while maintaining high mechanical specificity. Although frequency-domain and impulsive SBS were previously used for Brillouin imaging of tissue phantoms at high spectral resolution^21–23^, only a recent demonstration has achieved the necessary high spatial resolution and shot-noise sensitivity adequate for imaging of semi-transparent organisms, such as *C. elegans*^19^. However, this implementation used continuous wave (CW) laser illumination, therefore not taking full advantage of the non-linearity of the stimulated process. Moreover, the required high illumination dosages (~265 mW in Ref.^19^) limit their use for live-imaging of sensitive samples over extended time-periods which likely prohibits more widespread applications and its uptake in the life sciences.

In our work, we address this major drawback of current SBS realizations by introducing a quasi-pulsed pump-probe approach. Combined with balanced detection and diligent parameter optimization, this allows us to achieve shot-noise limited performance yielding comparable SNR, spectral resolution, image quality and speed compared to recent SBS demonstrations^19^, yet at a significantly (>10-fold) lower illumination dosages. We further show that, in contrast to full-CW operation, pulsed-SBS does not lead to sample heating or phototoxic effects in cultured single cells or during 2D live-imaging of entire organoids and *C. elegans* embryos over extended field-of-views (FOVs, up to 215×215×10 μm^3^) and time periods (up to 3 hours). The high spatial (0.57×0.55×2.58 μm^3^) as well as spectral (151 MHz) resolution and thus mechanical specificity further allowed us to distinguish up to three mechanically separate tissues in the focal volume in the zebrafish notochord and blood vessels, and to capture visco-elastic changes during organoid development.

## Results

### Pulsed SBS approach, setup and validation

The pulsed SBS approach and the schematic of our microscope is conceptually shown in **Fig. 1a-d**. It is based on two continuous wave (CW), narrowband (100 kHz), yet tunable, amplified 780 nm diode lasers that illuminate the sample from two opposing sides through high (0.7) numerical aperture (NA) objective lenses (**Online Methods**). The resulting interference pattern is determined by the spatial overlap of the pump and probe beams, which in our case are confined to 0.57×0.55×2.58 μm^3^ (see **Fig. 1d, SI Fig. 1**).

**Figure 1.**
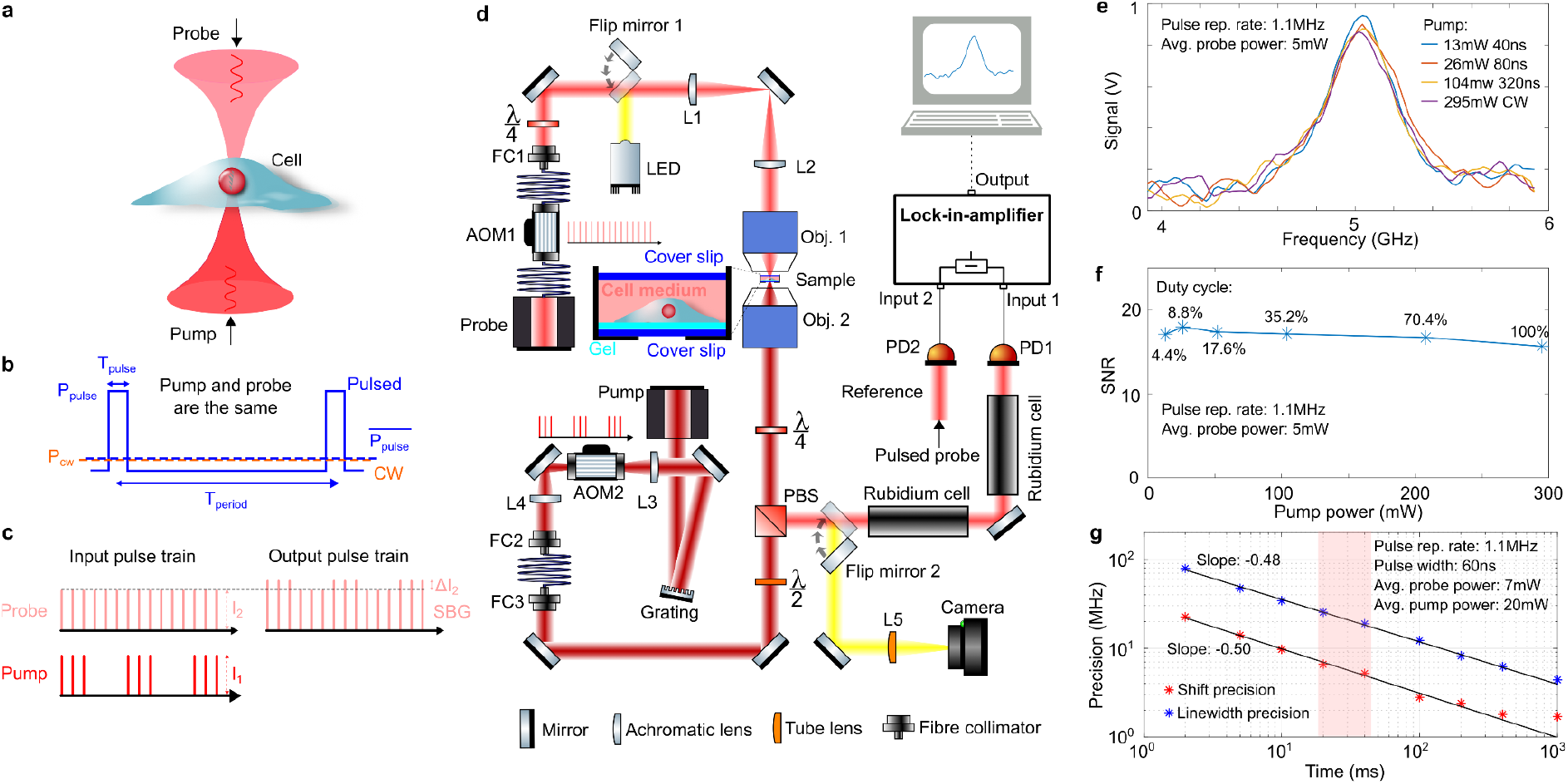
Pulsed-SBS approach and performance. **(a)** Pump and probe beams with a slightly different frequency counter-propagate and are focused inside a biological sample. **(b)** Schematic of the optical power against time for the quasi-pulsed scheme (blue solid line) and the CW scheme (orange dashed line). (**c**) Stimulated Brillouin gain (SBG) detection scheme. The pump beam is modulated at high frequency (320 kHz) at which the resulting amplitude modulation of the probe beam due to SBG can be measured. (**d**) Schematic of the SBS setup. The pump and probe beams are focused in the same position using high NA (0.7) objective lenses. The intensity of the probe beam is measured by a photodiode, connected to the lock-in amplifier. Brightfield imaging can be performed by flipping two mirrors into the optical path (yellow). Widefield fluorescence image is obtained by adding an excitation filter after the LED and an emission filter before the camera (not shown). (**e**) Pulsed-SBS spectra of water under different pulse width but at the same peak power (thus different average power) while keeping the same average probe power to 5 mW. The pump pulse width ranges from 40 ns to CW and the corresponding average power ranges from 13 to 295 mW. Integration time is 20 ms. The signal SNR stays constant in pulsed-SBS despite the lower average power, as quantified in (**f**). (**g**) Precision of the Brillouin shift and linewidth as a function of the integration time of the SBG spectrum of water. The precision is calculated as the standard deviation of the Brillouin shift and linewidth determined from the Lorentzian fits of n=300 SBG spectra measured sequentially. The slope close to −0.5 shows that the measurements are white noise limited within the measured range.

To illustrate the advantage of our pulsed SBS approach, consider the CW pump and probe power to have equal average optical power *P_cw_*, while the pulsed pump and probe both feature a peak power of *P_pulse_*, but a comparable average power of *P_cw_* (see **Fig. 1b**). Since the SBS signal is proportional to the pump and the probe power as well as their interaction time, for the CW scheme the signal is proportional to *P_cw_* * *P_cw_* * *T_period_* = 1 a.u., where *T_period_* is the pump-probe interaction time of the CW scheme. On the other hand, for the pulsed scheme the signal is proportional to 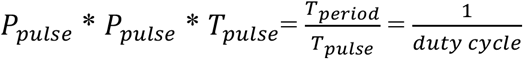 a.u., with *T_pulse_* being the pulse width. Therefore, the pulsed scheme yields an effective enhancement factor *E* that scales inversely with the duty cycle (*dc*), or *E* ~ 1/*dc* compared to the CW scheme with the same averaged pump and probe power. As the noise depends on the average probe power (which does not change), the pulsed scheme has *E* times higher SNR than the CW scheme. This enhancement can be employed to decrease the pump power while keeping the same SNR and probe power.

Quasi-pulsing of both the pump and probe lasers is achieved via acousto-optic modulators (AOM), which can be operated down to 40 ns length and 1.1 MHz repetition rate. In our work, we chose between 40 – 909 ns which corresponds to a *dc* of 4.4% - 100% or an averaged pump power dosage of 13 – 295 mW, respectively, and to a maximal *E*~22.7 (**SI Note 1**). For example, a pulsed scheme with 40 ns length for both pump (13 mW average power) and probe (5 mW average power) at 1.1 MHz repetition rate, has the same SNR as a CW-based scheme with a 295 mW pump and a 5 mW probe (see **Fig. 1e, f**). To record a Brillouin spectrum, the probe laser’s frequency is rapidly tuned across the Brillouin resonance of the medium, and the scattering signal is detected as an increase, or gain, of the probe’s intensity (I2). The typical spectrum acquisition time is as low as 20 ms in biological samples in this work. The detection unit comprises two photoreceivers for detection and reference respectively, and a differential-input lock-in amplifier (LIA), whose operation parameters were carefully optimized (see **SI Note 2, SI Fig. 2** and **Online Methods**). The sample is then raster-scanned through the microscope’s focus to obtain cross-sectional images of Brillouin shift (*Ω*_B_), linewidth (*Γ_B_*) and gain (*G*_*B*_).

First, we validated the pulse enhancement scheme by measuring the Brillouin water spectra and the signal-to-noise ratio (SNR) at different pulse length (hence different average power and duty cycle) and confirmed that SNR remains unchanged (see **Fig. 1e and 1f**). Furthermore, the precision of the Brillouin shift and Brillouin linewidth measurements which determine the Brillouin image quality, as a function of the acquisition time of the SBG spectrum of water are on par with recent SBS demonstration^19^ (**Fig. 1g**). It should be emphasized that with 20 ms integration time and 2 GHz frequency scan range, our pulsed setup with 20 mW average pump and 7 mW average probe (used in our Brillouin imaging experiments), has a shift precision of 6.6 MHz which is comparable to previous work using a low-NA CW setup with 250 mW pump and 15 mW probe^19^. The pulsed SBS-microscope also shows high spectral resolution (151 MHz, **SI Fig. 3**) as well as close to shot-noise limited performance across a wide range of LIA dwell times (**SI Note 2**). This demonstrates the pulse enhancement of our SBS approach and suggests state-of-the-art performance at effectively 22-fold reduced illumination powers.

### Cellular imaging with pulsed-SBS free of photodamage

To translate the advancements of the pulsed-SBS scheme to biomechanical imaging, we first acquired Brillouin maps of NIH/3T3 mouse fibroblasts, primary human brain microvascular endothelial cells (HBMECs) and mouse embryonic stem cells (mESCs), and compared them to images obtained when using quasi-CW illumination. As shown in **Fig. 2a** and **SI Fig. 4**, Brillouin shift maps acquired with pulsed-SBS (27 mW total power) showed overall high image quality and good SNR level (e.g. on average 32 dB in the first z-plane of Fig. 2a), from which subcellular compartments with subtle mechanical differences such as the nucleoli or nuclear envelope could be clearly discerned. From the spectral data, spatial maps of the Brillouin shift, gain and linewidth (**Fig. 2e, g, h**) both lateral as well as axial (**Fig. 2f, i**) can be plotted, demonstrating the high overall image quality and signal SNR. Staining with an established marker for cell death (propidium iodide, PI) revealed membrane and/or chromosomal damage was absent in a pulsed-SBS imaged cell 15 hours after imaging (**Fig. 2a**, n=5). Next, we sequentially imaged two fibroblast cells with pulsed and CW schemes. No cell blebbing and membrane damage were observed in the pulsed scheme (**Fig. 2b**). In contrast, while a Brillouin map acquired with SBS CW-illumination (250 mW total power) gave similar contrast and quality, it showed direct effects of potential photodamage: Firstly, clear cell blebbing and membrane damage were observed in the CW scheme after imaging (**Fig. 2c**). Secondly, the overall Brillouin shift (*Ω*_B_) was elevated in different cell regions (nucleolus, nucleoplasm and cytoplasm). This trend was observed in both fibroblast (n=2) and primary human brain microvascular endothelial cells (n=2), which suggests sample heating (**Fig. 2j**). Furthermore, mouse embryonic stem cells showed increased motility which is indicative of cellular avoidance behaviour (**SI Fig. 4e**). Together, these experiments emphasize that in SBS-based imaging of single cells, total illumination levels have to be carefully kept at low powers (<30 mW) to ensure cell viability. Here, pulsed-SBS is capable of retaining high image quality and mechanical sensitivity, opening the door for high specificity imaging of sub-cellular mechanics across various cell types and established cell lines.

**Figure 2.**
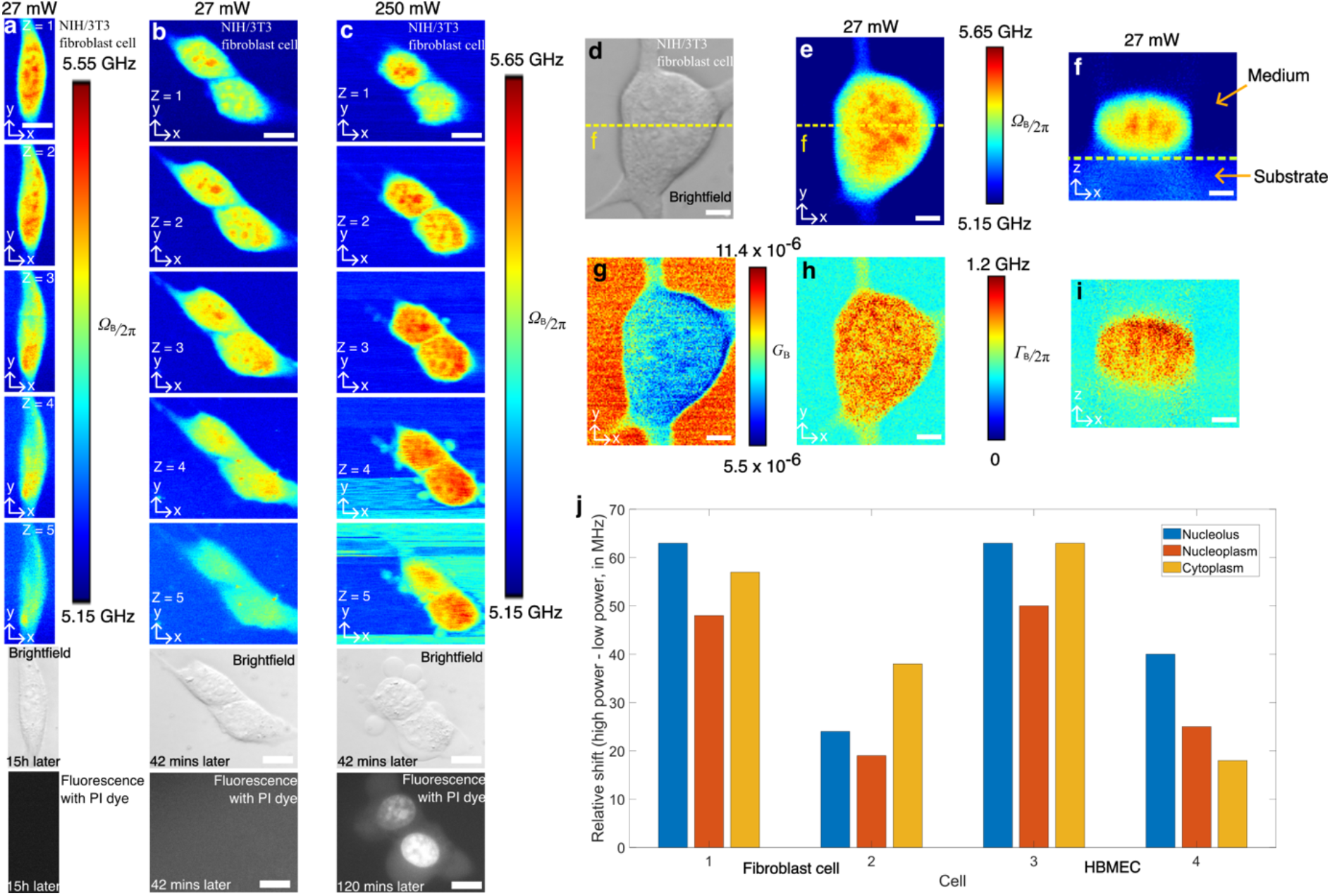
Pulsed-SBS imaging of cultured cells. **a**, Three-dimensional (3D) sections of Brillouin shift images of a NIH/3T3 fibroblast cell under 20 mW pump and 7 mW probe with a z-step of 1 μm. The coregistered brightfield image and fluorescence image (15 hours after Brillouin imaging) of the cell stained with propidium iodide (PI) dye are shown below. **b**, 3D Brillouin shift images of two fibroblast cells under 20 mW pump power and 7 mW probe power with a z step of 1 μm. The brightfield and fluorescence images (42 minutes after Brillouin imaging) are shown below the Brillouin images. **c**, 3D Brillouin shift images of the same two cells under 240 mW pump power and 7 mW probe power after (**b**) with a z step of 1 μm. Scale bars in (a)-(c) are 10 μm. Note that the PI dye is staining the nucleus, a clear indication of cell damage. **d**, Brightfield image of a NIH/3T3 fibroblast cell. **e**, **g** and **h** are respectively the Brillouin shift (*Ω*_B_), Brillouin gain (*G_B_*) and Brillouin linewidth (*Γ*_B_) images of the x-y plane. **f** and **i** are the Brillouin shift and Brillouin linewidth images of the x-z plane along the dashed yellow line in **d**. The dashed line in **f** marks the substrate (i.e. polyacrylamide-based gel) boundary. Scale bars in (d)-(i) are 5 μm. **j**, Relative frequency up-shift of CW-SBS with respect to pulsed-SBS of three regions (nucleolus, nucleoplasm and cytoplasm) for (n=4) cells between 27 mW power and 250 mW power. All image pixel steps and pixel time are 0.25 μm × 0.25 μm and 20 ms respectively.

### High specificity SBS imaging in live zebrafish and C. elegans adults

Next, we set out to explore the high spectral resolution of our pulsed-SBS for distinguishing different mechanical constituents in heterogeneous living tissues. For this we acquired low-power, pulsed-SBS images in the tail region of live zebrafish larvae 3 days post-fertilization (3dpf). Here, the region surrounding the notochord is known to encompass an ultra-thin, ~500nm wide, extra-cellular matrix layer of high mechanical rigidity that supports the tissue structure^13^. The notochord is an embryonic midline structure that serves as the axial skeleton and provides structural support to the developing embryo. Scanning a 60×20×32 μm^3^ FOV in the lateral and axial direction, centered ~500 μm anterior to the posterior end of the notochord, and plotting a single Brillouin shift peak location, revealed several tissue types including the notochord and the surrounding muscle segments including blood vessels (**Fig. 3c, d**). Owing to the high spectral resolution of SBS, an asymmetric spectrum indicative of several peaks becomes evident in several tissue regions. To aid in the efficient experimental evaluation and data analysis of the complex, multipeak Brillouin spectra, we developed a custom analysis approach and GPU-enhanced fitting pipeline with a convenient graphical interface and significantly improved speed which enabled real-time analysis and optimization of experimental parameters (**Online Methods, SI Fig. 5** and **Supplementary Software**). This allowed us to unambiguously distinguish between single and multiple peaks in our spectra (**Fig. 3e**). Using our optimized tools, we observe a clear triple peak distribution (**Fig. 3f**) of Brillouin shifts when analyzing the ECM region indicating that the ECM region consists of three mechanically distinct components. We interpret these as an ECM-surrounding tissue component (5.34 GHz, L1) which has a similar Brillouin shift as the nearby tissue, and the first (5.63 GHz, L2) and second (6.63 GHz, L3) ECM acoustic mode, as first described in purified ECM as bulk and parallel-to-surface modes^14^. Note that previous work^13^ in the zebrafish notochord was only able to distinguish the higher-shift, and thus well-separated, second ECM mode. Pulsed-SBS is therefore capable of identifying an unprecedented three distinct and specific biomechanical components inside a diffraction-limited focal volume and inside a living specimen. We also observed clearly double-peaked Brillouin spectra when analyzing presumable blood vessel regions (**Fig. 3g**), which we interpret as blood plasma (5.03 GHz, L1) and red blood cells (5.39 GHz, L2), in agreement with a similar report in the literature^24^. However, in contrast to those being derived from statistical analysis of multiple data points in Ref.^24^, we highlight that here we are able to measure both components inside the PSF simultaneously, i.e. for every image pixel.

**Figure 3.**
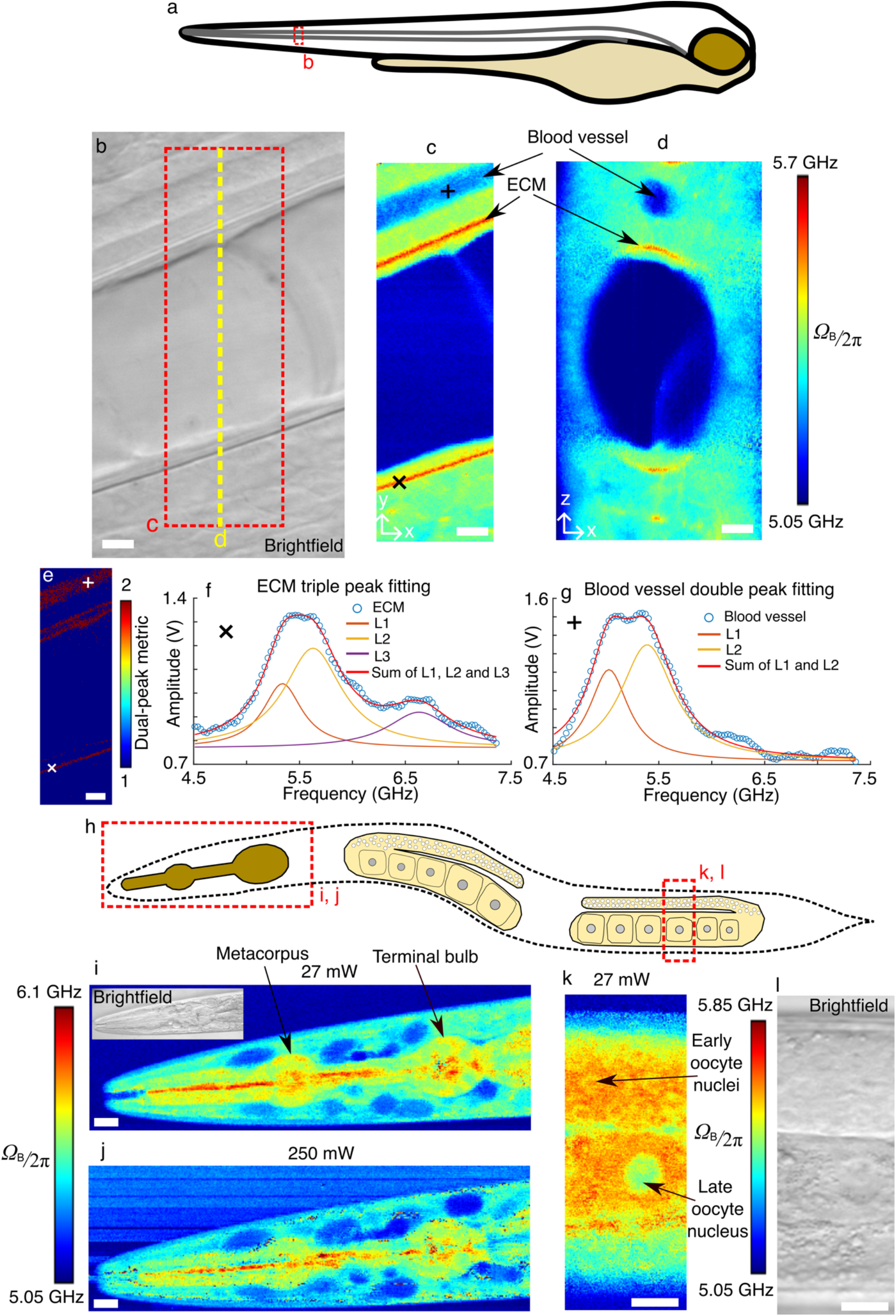
Pulsed-SBS imaging of zebrafish larvae and young adult *C. elegans*. **a**, Illustration of a zebrafish larvae, indicating the imaging region (dashed red box). **b**, Brightfield image of the notochord region of a zebrafish larva at 3 dpf stage. **c**, Brillouin shift image of the x-y plane of the region marked by a red dashed box in **b**. **d**, Cross-sectional Brillouin shift image in the x-z plane along the yellow dashed line in b. Scale bars in (b)-(e) are 5 μm. In **c** and **d**, the pixel steps in x and y directions are both 0.2 μm and the pixel time is 40 ms. **e**, Multi-peak metric map of the region marked by a red dashed box in **b**. **f**, A representative Brillouin spectrum in the high-shift region, indicative of extracellular matrix (ECM), marked by a cross in **c** and **e**. The raw spectrum and its triple-peak fitting are shown, revealing distinct mechanical components (L1-3). Note that the offset was added to L1-3 for visualization. **g**, A representative Brillouin spectrum in the blood vessel region marked by a plus sign in **c** and **e**. The raw spectrum and its double-peak fitting (L1, L2) are shown. Note that the offset was added to L1 and L2 for visualization. **h**, Schematics of a young adult *C. elegans*. The head and gonad regions imaged are marked by dashed red boxes. Brillouin shift images of the head region under 27 mW power (**i**) and 250 mW power (**j**). In **i** and **j**, the pixel steps in x and y directions are both 0.5 μm and the pixel time is 20 ms. Brillouin shift image (**k**) and coregistered brightfield image (**l**) of the gonad region marked by a dashed red box in **h**, showing oocyte nuclei from different meiotic stages (arrows). In **k**, the pixel steps in x and y directions are both 0.25 μm and the pixel time is 20 ms. Scale bars in (**i**)-(**l**) are 10 μm.

We further validated the quality of our pulsed-SBS microscopy by imaging live, anesthetized wild-type *C. elegans* at the young adult stage. As evident from close-up images of the pharyngeal region (**Fig. 3i, j**), pulsed-SBS yields similar SNR (e.g. on average 23dB in **Fig. 3i**) and overall contrast compared to standard, CW-SBS^19^ at comparable acquisition parameters (3 GHz scan range, 20 ms spectral acquisition time), yet at 10-fold reduced illumination power. Here, under CW-illumination, optical trapping of dust in the illumination laser light focus, particularly in the media and lumen, were more common and the specimens imaged showed enhanced motility behaviour despite anesthesia. This behavior is likely the response of the worm to prolonged exposure of high intensity near infrared light which can also trigger a stress response in the animal^25^. We therefore used our pulsed-SBS to observe spatial differences in Brillouin shift, and hence stiffness, in the gonad region of the nematode at high resolution (**Fig. 3k**). The stiffness of meiotic nuclei decreased during oocyte maturation. While a thin shell of softer nucleoplasm was wrapped around a stiff nucleolus in early oocyte nuclei, both nucleoplasm and nucleolus regions were very soft in late oocytes. This decrease in stiffness correlates with an increase in chromatin compaction from early to late oocytes.

To further highlight the advantage of low-power pulsed-SBS over standard CW-SBS approaches for imaging sensitive biological specimens, we also imaged live mouse embryos in the zygote and 8-cell-stage (**SI Fig. 6**). We also obtained high image quality and sub-cellular details with pulsed-SBS, while the high optical power of CW-SBS trapped the embryo and thus prohibited Brillouin imaging. We note that the lack of proper environmental control (temperature, CO_2_, etc.) prevented studying mouse embryos over longer time periods, which will be the subject of future studies.

### Visco-elastic imaging of mouse mammary gland organoids

To demonstrate the applicability of our method for medically relevant test systems, we imaged organoids, which are seen as emerging models to investigate healthy and diseased tissues. Here, we acquired pulsed-SBS images of mouse mammary epithelial organoids (**Fig. 4**). The epithelial tissues and cells were well resolved in lateral as well as axial cross-sections. Quantifying the Brillouin linewidth (*Γ_B_*) we observed a transient decrease in linewidth (~190 MHz) and thus viscosity in the organoid lumen between early (48 hours post-seeding the single cells into basement membrane matrix) (**Fig. 4b, c**) and later phases (70 hours and 80 hours post-seeding) (**Fig. 4e-k**) of development. In this organoid system, lumen formation is generated through cell division^26^ and hollowing^27^, that typically involves endocytosis of membrane vesicles to extend the apical membrane^28^ as well as hydrostatic pressure, which builds through the accumulation of ion channels and pumps at the apical plasma membrane^29,30^. These processes may explain the observed increased viscosity in the early phase of organoid growth, when a nascent lumen is formed (**Fig. 4b, c**). We furthermore captured a putative cell division event during late cytokinesis (**Fig. 4d**). Remarkably, these cells showed a significant decrease in Brillouin shift (see arrows in **Fig. 4e, f**), very much in line with observations in the Madin-Darby Canine Kidney (MDCK) cell line at the division elongation/cytokinesis stage in 2D cell culture conditions as measured by traction-force microscopy^31^.

**Figure 4.**
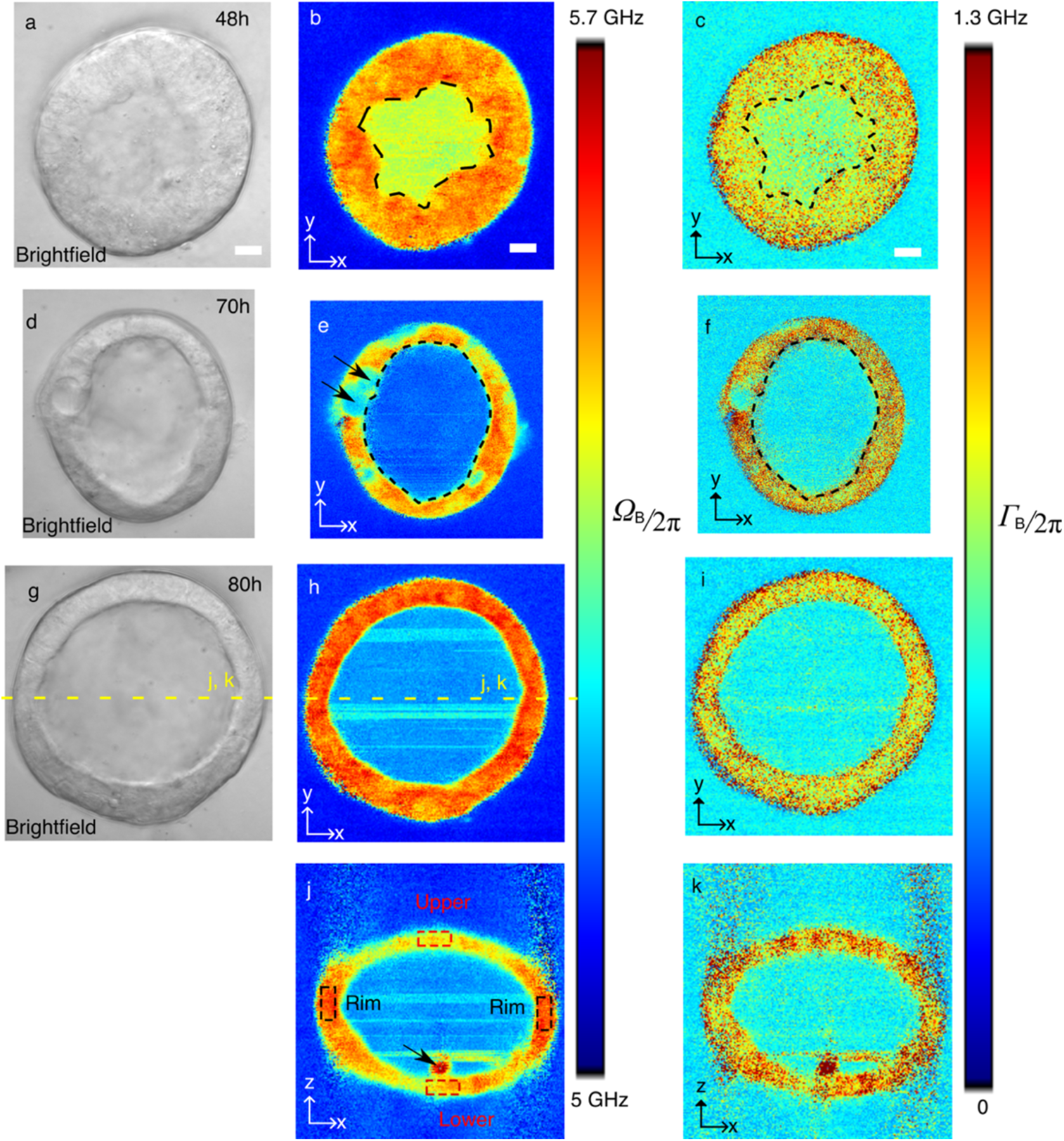
Pulsed-SBS imaging of mouse mammary gland organoids. **a**, Brightfield image of an organoid 48h post seeding singularized mammary epithelial cells into basement membrane matrix. (**b**) Brillouin shift image and (**c**) Brillouin linewidth image of the organoid in **a**. In **b** and **c**, the pixel steps in x and y directions are both 0.5 μm and the pixel time is 20 ms. **d**, Brightfield image of an organoid 70 h post seeding. (**e**) Brillouin shift image and (**f**) Brillouin linewidth image of the organoid in **d**. The average Brillouin linewidth in the lumen in **f** (FWHM 488 MHz) is 200 MHz decreased compared to **c** (FWHM 688 MHz) indicating lower viscosity in older organoids. Arrows in **e** highlight a dividing cell in cytokinesis. Dashed lines in b-f demarcate organoid lumen. In **e** and **f**, the pixel steps in x and y directions are both 0.25 μm and the pixel time is 20 ms. **g**, Brightfield image of an organoid 80h post seeding. (**h**) Brillouin shift image and (**i**) Brillouin linewidth image of the organoid in **g**. (**j**) Brillouin shift image and (**k**) Brillouin linewidth image of the x-z plane marked with a dashed yellow line in **g** and **h**. The average Brillouin shift of the bended rim regions (black dashed boxes in (j), average 5.56 GHz) shows a ~100 MHz up-shift compared to the average shift of the upper and lower cap of the epithelium (red dashed boxes in (j), average 5.46 GHz). Note that even with only 27 mW power, particle trapping leads to stripe-like artefacts in the lumen of the organoids. A piece of debris, potentially a dead, extruded cell is marked with an arrow in j. In **h**-**k**, the pixel steps in x and y directions are 0.5 μm and the pixel time 20 ms. Scale bars in all the images are 10 μm.

Finally, in 80-hour old, established organoids, we observed differential Brillouin shift and hence mechanical rigidity in axial cross-sectional images, with the side epithelium displaying a 100 MHz higher shift (**Fig. 4j, k**). Overall, organoids remained viable after pulsed-SBS imaging, as confirmed by brightfield microscopy acquired at 24 and 48 hours after the imaging experiments, that showed intact organoids and no detectable blebbing or debris originating from dead cells within the lumen.

### Pulsed-SBS enables live-imaging of C. elegans embryo development

As a final demonstration of the low-phototoxicity of our pulsed-SBS method and its capability to visualize dynamic tissue properties throughout morphogenesis, we captured longitudinal and 2D mechanical information of the developing *C. elegans* embryo, from ~300-480 minutes post-fertlization (mpf), i.e. from “bean” stage to “1.5 fold” stage (**Fig. 5** and **SI Video 1**). This is the period of embryonic development when tissue morphogenesis starts, after the majority of embryonic cell divisions have concluded^32^. We acquired 2D Brillouin time-lapse images over a ~65×55 μm^2^ FOV within ~4.8 min and at 13 min time-intervals (i.e. over 15 separate timepoints). Analyzing our Brillouin time-lapse data, we observed differential Brillouin shifts, and hence stiffnesses, across embryonic body parts and embryonic stages. During early morphogenesis, higher stiffnesses mark the posterior part of the embryo mostly consisting of endoderm cells which will form the intestine, in contrast to the anterior embryo part mostly composed of nervous cells (**Fig. 5a-h**). Later, at 400-450 mpf (**Fig. 5h-o**), stiffness values remain high close to the intestine, while also increasing within the anterior of the embryo where the formation of the major brain neuropil and head muscles are being established^32,33^. No photodamage or -toxicity was observed at ~27 mW of average laser power as confirmed by brightfield images of the embryos post-acquisition (n=3 embryos). *C. elegans* embryos remained active and viable after pulsed-SBS imaging. Brightfield imaging acquired at 10 hours after the last Brillouin acquisition time (~440 mpf), confirms that the imaged embryos undergo twitching and normal development past the stage of “4-fold” which precedes the hatching of embryos into larvae animals. Moreover, 3 out of 3 embryos imaged with pulsed-SBS imaging all eventually hatched into viable larvae. This represents a significant improvement over standard CW-SBS employing ~265 mW, where embryo damage and death was observed following the acquisition of a few 2D images (n=3 embryos, **SI Fig. 7**).

**Figure 5.**
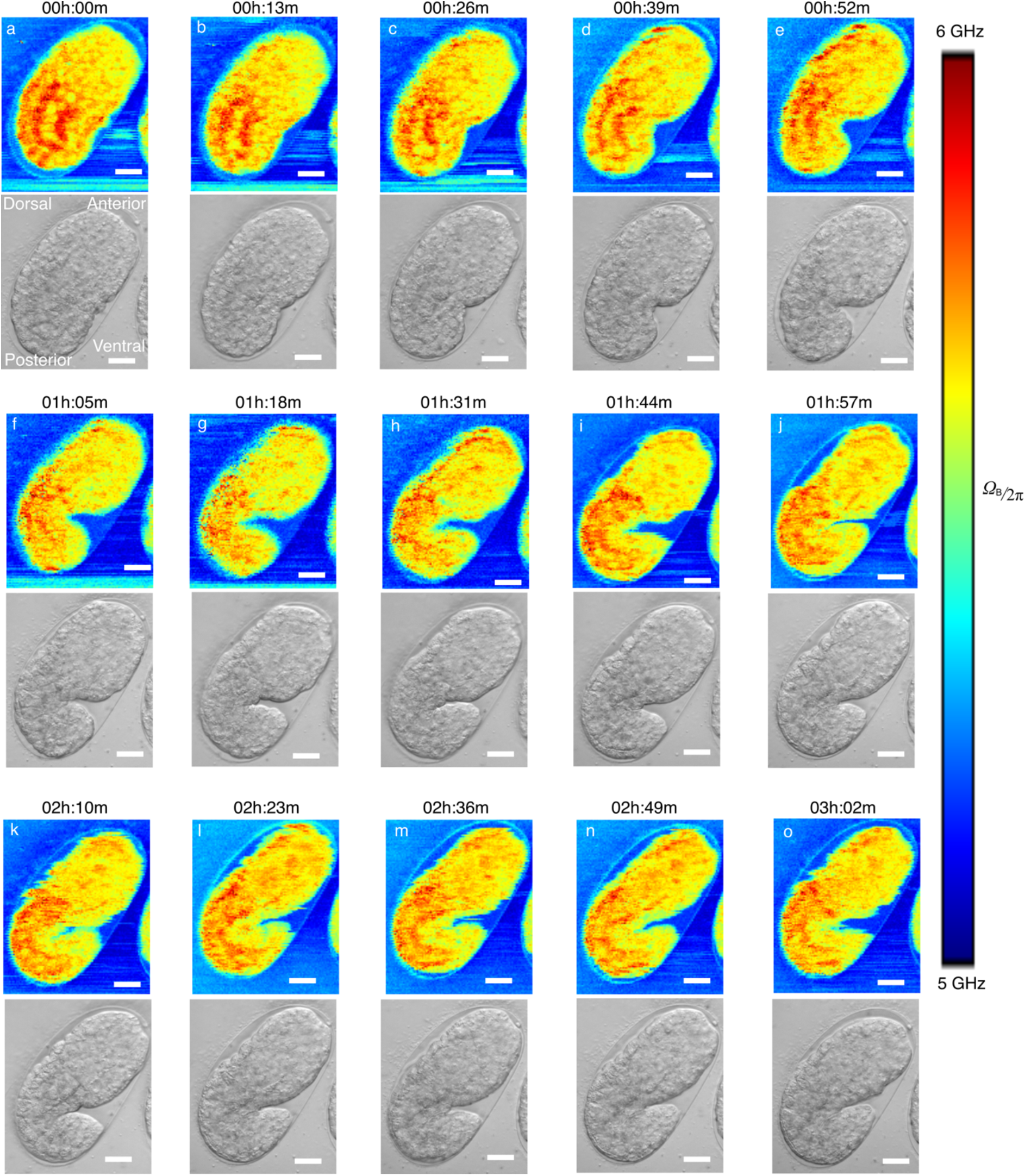
Time-lapse pulsed-SBS imaging of *C. elegans* embryo development. Brillouin shift (top) and brightfield (bottom) images over a 3-hour time-span and at 13min time interval. The pixel steps in x and y directions are 0.5 μm and the pixel time is 20 ms (total time per image 288 sec). Scale bars in all the images are 10 μm. The embryo at the start of the time-lapse is at the late “ball-stage”, slightly before the bean stage, or approximately at ~300 minutes post-fertilization (mpf). Imaging timelapses presented here conclude at ~480 mpf, when embryo start their first spontaneous muscle contractions, known as “twitching”.

## Discussion

To summarize, here we presented a new scheme for stimulated Brillouin microscopy that fully exploits the non-linearity of the pump-probe interaction. In our work we harnessed the improved efficiency of our approach along with diligent optimization of the signal detection to substantially reduce the required total illumination power and thus enable imaging of a wide range of photosensitive biological samples previously inaccessible to SBS. These advances have allowed us to demonstrate non-invasive 3D live-imaging of various cell types, developing embryos and organisms as well as tissue organoids. Moreover, the high mechanical specificity of our method together with an enhanced data analysis pipeline facilitates the distinguishing of different mechanical constituents within diffraction-limited volumes inside heterogeneous living bio-samples.

Our pulsed SBS approach achieves ~20-fold signal enhancement at a noise level only two-fold higher than the shot noise. Therefore, 10-fold lower power illumination could be employed to achieve the same SNR as the most recent state-of-the-art previous work^19^. Ensuring low photo-burden is of utmost importance when establishing new imaging and spectroscopy methods for useful applications in biomedicine. Here we note that our 10-fold / 20dB power reduction (because of 20 dB signal enhancement) compared to the shot noise limited CW scheme is significantly higher than recently achieved quantum-enhanced spectroscopy schemes for Raman^34^ (35% SNR enhancement) and SBS^35^ (3.5 dB SNR enhancement).

While the spectral and thus overall image acquisition time of pulsed-SBS (~20 ms) is notably slower than recently developed line-scan Brillouin microscopy approaches^36–38^, we note that it is still faster than standard confocal implementations^8,12,39,40^, yet yields improved spectral linewidth fidelity and thus mechanical specificity, and in principle allows information to be obtained on the mass density. The limitations of pulsed-SBS are that it is presently restricted to relatively (i.e., non-absorbing) samples thinner than ~100-200 μm that can be optically accessed from two opposing sides. Furthermore, at present, the sample is raster-scanned under the microscope, which for very fast scans and/or large step sizes can lead to artifacts from sample movement, and the input power of the fiber-pigtailed probe AOM is limited to ~650 mW, thus limiting the overall duty cycle and enhancement factor to ~20 (**SI Note 1**) with a reasonably good SNR.

Future hardware developments will allow us to fully exploit the possible enhancement of our pulsed-SBS approach. In particular, nanosecond pulsed laser light generated by faster electro-optic modulator (EOM), together with smaller repetition rate (e.g. 100 kHz), could unlock the remaining untapped potential. Here, we envision that diligent illumination scheme design could achieve overall enhancements of up to ~10.000-fold, which could be translated to practical spectral acquisition times of as low as ~20 μs at <1mW average illumination power only (**SI Note 1**). Such radical improvements could open new avenues and further applications for studying biomechanics in 3D with high spatio-temporal resolution and mechanical specificity in living biological specimen in a truly non-invasive manner, providing transformative capabilities for the field of mechanobiology.

## ACKNOWLEDGMENTS

We would like to thank the mechanical and electronic workshop at EMBL Heidelberg for help; A. Bilenca for helpful advice during early stages of the project; M. Schindler and N. Petridou for providing zebrafish larvae. A.D.-M. was supported by the Deutsche Forschungsgemeinschaft (DFG) research grants DI 2205/2-1 and DI 2205/3-1. R.P. acknowledges support of an ERC Consolidator Grant (no. 864027, Brillouin4Life), the German Center for Lung Research (DZL), research funding “Life Science” of the Molit Institute, and a COST Innovator grant (IG16124). R.P. and A. D.-M. acknowledge funding from the COST Action CA16124 (‘BioBrillouin’). This work was supported by the European Molecular Biology Laboratory.

## AUTHOR CONTRIBUTIONS

F. Y. and R.P. conceived the project. F.Y. and R.P. designed the imaging system, and F.Y. realized it with help from C.B. S.H. wrote the custom spectral analysis code. F.Y. performed experiments and analyzed data, with help from G.R., as well as from A.N, A.G., K.W and MG., under guidance of S.K., A.D.-M., J.E. and M.B., respectively, R.P. led the project and wrote the paper together with F.Y. and input from all authors.

## Online methods

### SBS theory

Instead of employing the interaction between single photons and phonons generated by thermal agitation in spontaneous Brillouin scattering, stimulated Brillouin scattering uses two light beams. Pump (*ω*_1_) and probe (*ω*_2_) beams with a slightly different frequency counter-propagate and are focused in the sample, sharing the same focal volume. The beating between the pump and probe generates an interference fringe pattern, whose contrast is time-varying at the frequency given by the difference between the pump and probe frequency. Due to electrostriction, the interference fringe modulates the density of the sample and hence generates an acoustic wave. Because of the photoelasticity effect, the acoustic wave changes the refractive index of the sample and generates a moving Bragg grating which can transfer the energy from the pump to the probe beam. Therefore, this results in a stimulated Brillouin gain (SBG) for the probe beam, while for the pump this represents a stimulated Brillouin loss (SBL) process. The process turns resonant when the beat pattern moves at the sound velocity in the sample, which is realized for a well-defined frequency difference (*Ω*_B_ = *ω*_1_-*ω*_2_) between the pump and probe beams.

The SBG or SBL spectrum is given by *G*(Ω) =±*η* × *g*(Ω) × *l* × *I*_1_, where *η* is the overlap efficiency of the pump and probe beams in the sample, ±*g*(Ω) is the SBS gain or loss factor well described by a Lorentzian function, *l* is the interaction length of the two counter propagating laser beams in the sampled volume, and *I*i is the intensity of the pump beam at *ω*_1_. As for the spontaneous Brillouin scattering spectrum, the close relationship between the SBG or SBL spectrum and the complex longitudinal modulus of the probed volume allows to locally extract the high-frequency viscoelastic response of the medium. This relationship is described by *M*\* = *ρ* × (*λ*_1_/2*n*)^2^ × Ω_B_^2^ × (1 +iΓ_B_/Ω_B_), where *M*\* is the complex longitudinal modulus, *λ*_1_ is the wavelength of the beam at *ω*_1_, and *n* and *ρ* are the refractive index and the mass density of the medium, respectively. Similar to other nonlinear optical techniques, SBS inherently provides optical sectioning in three dimensions owing to the nonlinearity of SBS in the total irradiance intensity.

### Pulsed SBS imaging setup

A detailed schematic of our pulsed-SBS microscope is shown in Fig. 1d. Two tapered-amplified CW external-cavity diode lasers (TA-Pro, Toptica) with linewidth of 100 kHz are used as pump and probe beams. The wavelength of the pump laser is kept at 780.24 nm which is at the D2, F=3 absorption line of Rb^85^. To get the SBG spectrum, the probe laser frequency is scanned across the Brillouin gain peak *Ω*_B_) by sawtooth modulation of the laser piezo. The probe beam (s-polarization) is amplitude modulated by a fiber-pigtailed acousto-optic modulator AOM1 (#TEM-250-780-2FP-HP, Brimrose) to generate a pulse train (e.g. 60 ns pulse width, 1.1 MHz repetition rate). An aspheric lens (#49-115, Edmund) is used to collimate the output of the AOM to a collimated beam with a diameter of 6.7 mm (1/e^2^, theoretical). The probe beam is then right circularly polarized by a quarter wave plate. Two mirrors, conjugated to the back and the front focal plane of the probe objective (Obj. 1) by means of two achromatic lenses (L1 and L2: #49-356, Edmund), respectively, allow for fine position and angle adjustment of the beam inside the sample in order to aid in the alignment between the pump and probe overlap.

For the pump beam, a holographic narrow-band reflective Bragg grating (SPC-780, OptiGrate) is employed to clean up the pump laser’s amplified spontaneous emission noise. The filtered pump beam with s-polarization is focused into a free-space AOM2 by an achromatic lens (AC254-100-B-ML, Thorlabs) and re-collimated by another achromatic lens (AC254-100-B-ML, Thorlabs). AOM2 is not only used for pulse generation (e.g. 60 ns pulse width, 1.1 MHz repetition rate) but also for envelope amplitude modulation at 320 kHz frequency so that the pump modulation can be transferred to the probe by SBG and be measured by a lock-in-amplifier (LIA, MFLI 5MHz, Zurich Instruments). A single-pole, single-throw analog switch (#NC7WB66K8X, Onsemi) is used to electronically combine the pulse and the envelope amplitude signals. The pulse signal (with 60 ns pulse width, 1.1 MHz repetition rate) is used as a digital enable input which allows the input signal (sinusoidal envelope modulation at 320 kHz) to pass (or not). The pulsed pump beam is coupled into a fiber by a fiber coupler FC2 (FiberDock, Toptica) for beam delivery and is then collimated with a diameter of 6.7 mm (1/e^2^, theoretical) by a fiber collimator (#49-115, Edmund). Next, the beam is p-polarized by a half wave plate and transmitted through a polarizing beam splitter (CCM1-PBS25-780M, Thorlabs). The polarization of the beam is then changed to left circular polarization by a quarter wave plate.

The right-circular polarized probe beam and left-circular polarized pump beam counter-propagate and are focused into a sample at the same position by two 0.7 NA objective lenses (LUCPLFLN60X, Olympus). Note that Obj. 2 is mounted on a 3D translational stage for optimization of the overlap between pump and probe beams. Three-dimensional images are obtained by raster-scanning the 3D piezo stage (L3S-D10300-XY300Z300, nanoFaktur) onto which the sample is mounted. After interacting with the pulsed pump beam, the SBG probe beam is collected by Obj. 2 and its polarization is changed from right-circular polarization to s-polarization by the quarter wave plate. The s-polarization SBG probe beam is then reflected by the polarization beam splitter and passes through two Rubidium cells (SC-RB85-(25×150-Q)-AR, Photonics Technologies) before being detected by a photodiode PD1 (FDS1010, Thorlabs). The output of PD1 is split to DC and AC components by a bias-tee (ZFBT-4R2GW+, MiniCircuits). The DC component is measured by a data acquisition card (DAQ, National Instruments) for gain calculation and balance detection. The AC signal is filtered by a low-pass filter (LPF-B0R35+, MiniCircuits) and then measured by the differential input #1 of the lock-in-amplifier (LIA). A fiber coupler takes out 50% of the pulsed probe beam for balance detection. The other 50% go through a variable ND attenuator and is detected by PD2 (FDS1010, Thorlabs). The output of PD2 is also splitted to DC and AC components by a bias-tee (ZFBT-4R2GW+, MiniCircuits). The DC component is measured by the DAQ card as a reference beam for balance detection. The AC signal is filtered by a low-pass filter (LPF-B0R35+, MiniCircuits) and then measured by the differential input #2 of the LIA. It should be emphasized that the LIA is set to differential input mode for balance detection which can drastically decrease the detection noise (see Supplementary Figure 1). The LIA demodulated signal is acquired by the DAQ card. The reflection of the pump beam and the stray pump light can be filtered out by two Rubidium cells.

To acquire bright-field and wide-field fluorescence images, flip mirror 1 is flipped down and flip mirror 2 is flipped up. A lamp (X-Cite 120Q, Excelitas) with a light guide and a coupling adapter (X-Cite Light Guides, Excelitas) is used for brightfield imaging and the adapter output is conjugated to the back focal plane of Obj. 1 for Köhler illumination. The brightfield image is recorded by a CMOS camera (CM3-U3-31S4M, Flir).

To acquire wide-field fluorescence images of the PI dye for cell viability tests, a green LED (M530L4, Thorlabs) is butt-coupled to a light guide with a coupling adapter (X-Cite Light Guides, Excelitas) and used as an illumination light source. An excitation bandpass filter (ET535/50m, Chroma) and detection long-pass filter (ET605LP, Chroma) are placed after the LED and before the tube lens, respectively, for fluorescence detection.

### Pulsed-SBS system characterization and image acquisition

#### Optimization of pump-probe overlap

To maximize the overlap efficiency *η* of the pump and probe beam as well as to ensure the best axial resolution, the correction collars of the two 0.7 NA objectives require fine-tuning. First, the overlap between the pump and probe beam is roughly optimized (i.e. pump and probe are focused in the same position in the sample) by maximizing the coupling probe power out of FC2 in Fig. 1d. Next, the correction collar of Obj. 2 is adjusted to make the brightfield image of a specimen (e.g. a cell) as sharp as possible. While doing so, the z position of the sample needs to be adjusted, iteratively, to keep it in focus. After this procedure the best aberration correction for Obj. 2 is achieved and Obj. 1 can be optimized by maximizing the overlap efficiency *η* as described before.

#### Spatial and spectral resolution measurements

The lateral (i.e. x and y) resolution is characterized by measuring a PDMS bead in 1% (w/v) agarose (see SI Fig. 1(a)-(d)). The Brillouin frequency shifts of the agarose and PDMS bead are 5.12 GHz and 4.07 GHz respectively. We plot the fitting signal amplitude at 5.12 GHz in SI Fig. 1(b). Then we fit the amplitude across the edge of the bead in x and y directions with erf function. Therefore, we get the x and y FWHM resolution of 0.57 μm and 0.55 μm, respectively.

Because of the large refractive index difference between PDMS bead (RI=1.43) and agarose (RI=1.33), the aberration is high in the region close to the axial boundary of PDMS bead and agarose. We characterize the axial resolution by measuring immersion oil sandwiched between two cover slips (see SI Fig. 1(e) and (f)). The Brillouin frequency shift of the oil is 7.04 GHz and the frequency shift of the cover glass is more than 15 GHz, thus the cover glass Brillouin signal is outside our scanning range (5-9 GHz). We scan the focus point through the boundary of oil and glass and the signal amplitude at 7.04 GHz is shown in SI Fig. 1(f). The z axial resolution is calculated to be 2.58 μm by fitting the erf function to the experimental data.

To obtain the spectral resolution of the pulsed-SBS microscope, double-distilled water SBG signal was measured at 23°C and representative spectrum is shown in Supplementary Fig. 3. The FWHM linewidth is 459.2 MHz for the integration time of 20 ms and LIA noise-equivalent bandwidth of 200 Hz. Therefore, the spectral resolution of the microscope is calculated to be 151 MHz by subtracting our measured water FWHM to the low NA water FWHM (308 MHz measured in Ref.^19^). Note that the water FWHM linewidth is measured to be 410 MHz (corresponding to 102 MHz spectral resolution) when the integration time is 40 ms and LIA noise-equivalent bandwidth is 200 Hz, which is in agreement with the measured high (0.7) NA spectral broadening of the CW scheme^19^.

#### Determination of signal SNR

In our pulsed-SBS microscope, there are three types of contrast: Brillouin frequency shift *Ω*_B_, Brillouin linewidth *Γ*_B_ and Brillouin gain *G*_B_. *Ω*_B_ and *Γ*_B_ of each pixel are respectively the peak frequency and FWHM linewidth of the Lorentzian fitting of the measured spectrum. *G*_B_ of each pixel is the ratio between the amplitude of the Lorentzian fitting and the DC component of the photodiode output.

The signal intensity is the amplitude of the Lorentzian fitting from the measured spectrum. The noise is the standard deviation of the measured spectrum when the pump-probe frequency difference is tuned away from the Brillouin frequency shift. For example, to get the water SBG spectrum, the frequency difference was scanned from 4 to 6 GHz. The signal intensity is the amplitude of the Lorentzian fitting. The noise is the standard deviation of the spectrum outside the Brillouin interaction (i.e. 7-9 GHz).

### SBS spectral data analysis

The data acquired from the microscope is stored as a unidimensional array. The data is reshaped into a 2D array of dimension [frequency;pixels], and a high-pass filter is applied (cut-off frequency 50 Hz, corresponding to the typical acquisition time of a single spectrum). The recorded data is then processed using a customized version of Gpufit^41^ to which a Lorentzian function and a sum of two Lorentzian functions were added as fitting functions. Each Lorentzian fit gives access to the shift, width and amplitude of the peak. To plot an image or an image stack from this data, the array is reshaped, every second line is shifted (by a number of pixels equivalent to ~100 ms) to compensate for the delay between desired and actual stage position and every other plane is flipped (Z-scan pattern).

Except for the modified version of Gpufit, which is written in C++ and Cuda, the rest of the processing is done in Matlab. This allowed the quick development of a GUI to explore the acquired data, to detect problems in the acquisition and to verify the goodness of the fits. This GUI displays 2 images made from a selected parameter (shift, width, amplitude, error, etc…), and clicking on a pixel in these images displays the underlying acquired spectra and the fitted function (see SI Fig. 5).

In some cases, it is possible that the recorded spectra is a “double peak”, i.e. the sum of 2 Lorentzian functions. In that case, Gpufit is used to directly fit the sum of 2 Lorentzian functions. To determine if a given pixel is a single or a double peak, we evaluate the Akaike Information Criterion (AIC) to test for the appropriateness of a multi-peak fit. However, we found that this test alone is not sufficient as the recorded signal is not always a perfect symmetrical Lorentzian^42^.

To distinguish potentially asymmetrical peaks from true double peaks, we compute the derivative of the fitted sum of Lorentzian functions, and evaluate whether it intersects more than once with the y=0 line, in which case it is indicative of at least 2 local extrema. Furthermore, we slightly shift the derivative offset and repeat the test. In case we also get intersections with y=0, it points to one local extrema in the spectrum and a ‘bump’ on the side. Both these situations are clearly impossible to have with only one, either symmetrical or asymmetrical, peak. We found that shifting the derivative offset from −0.5 to +0.5 allowed us to classify 99.9% of pure water spectra as a single peak.

On our workstation (11th Gen Intel(R) Core(TM) i9-11900K @ 3.50GHz and Nvidia GeForce GTX 1050 Ti), using the GPU and Gpufit instead of using the default Matlab fitting function enables a speed-up of the total processing time of ~340 times (see SI Table 1).

### Sample preparation and imaging parameters

#### *Cells* (incl. staining and controls)

Mouse fibroblasts (NIH/3T3-CRL-1658 from ATCC) were grown in media composed of DMEM with 4.5g/L glucose, 10% fetal bovine serum (Gibco, Cat.# 26140079) and 100U/mL penicillin-streptomycin (Gibco, Cat.# 15140122). They were seeded on polyacrylamide (PAA) gels with 12 kPa hydrogel stiffness (# SV3510-EC-12, Matrigen) at a density of 4000 cells/cm^2^ and allowed to adhere overnight before imaging.

Mouse embryonic stem cells (sox1-GFP-mESCs from Austin Smith’s lab, University of Exeter) were maintained as undifferentiated stem cells in media composed of high glucose DMEM (Gibco, Cat.# 11960044) supplemented with 15% ES-qualified fetal bovine serum (Merck, Cat.# ES009B), 1x NEAA (Gibco, Cat.# 11140035), 1mM sodium pyruvate (Gibco, Cat.# 11360039), 0.1mM beta-mercaptoethanol (Gibco, Cat.# 21985023), 1 U/ml Leukemia Inhibitory factor (from EMBL protein expression and purification core facility), 2mM L-glutamine (Sigma, Cat.# G7513) and 100U/ml penicillin-streptomycin. Media was refreshed every day and the cells were subcultured every second day with a seeding density of 10000 cells/cm^2^. PAA gels were coated with 0.1% Bovine skin Gelatin type B (Sigma, Cat.# G9391) in phosphate-buffered saline (PBS) for 1 hour at 37°C prior to seeding mESCs at a density of 10000 cells/cm^2^. The cells were allowed to adhere overnight before imaging.

Primary human brain microvascular endothelial cells (HBMECs, Cell systems), kept to a passage number lower than 10, were cultured in EGM-2 MV media (Lonza) on flasks coated with poly-l-lysine (Sigma Aldrich). HBMECs were then seeded onto dishes with 30μl of 7.5mg/ml collagen I solution (isolated from rat tails) spread evenly. HBMECs were allowed to attach and spread on the collagen overnight before being imaged the next day.

To test for cell viability under low power pulsed-SBS and high power CW-SBS, we added propidium iodide (≥ 94% PI, P4170-10MG, Sigma-Aldrich) into the cell dishes with a volume ratio to the cell medium of 30 μL in 2 mL.

#### Zebrafish

A wild-type zebrafish larvae (3 dpf) was used in Fig. 3a-g. 1-pheny1 2-thiourea (PTU) was added at 0.003% concentration at 12 hours post fertilization to avoid pigmentation. The fish was mounted in the center of a petri dish (Mattek, 35 mm Dish, No. 0 Coverslip) with 1% (w/v) low melting agarose and 0.016% (w/v) tricaine (for immobilization purpose). A coverslip was used to cover the agarose before solidifying to allow the access of the sample by two objectives.

#### C. elegans

##### *C. elegans* adult preparation

N2 Bristol animals, 6h post-L4, were washed twice in M9 buffer (22 mM KH_2_PO_4_, 49 mM Na_2_HPO_4_, 86 mM NaCl, 10 mM NH4Cl) with 0.1% (v/v) Tween-20 followed by two washes in M9 buffer with 0.1% (w/v) tetramisole (mounting solution). The animals were mounted in 5 μL of mounting solution between two agarose pads of 120-μm thickness each. To make agarose pads, 18 μL freshly prepared 5% (w/v) low melting agarose in M9 buffer with 0.01% (w/v) tetramisole were pipetted onto a glass slide and pressed against another slide. To control for the agarose pad thickness, a 120-μm spacer was placed on the slide containing the agarose. The final sample thickness was ~240 μm.

##### *C. elegans* embryo preparation

*C. elegans* adults of the N2 Bristol strain were isolated on agar plates and allowed to lay eggs, corresponding to embryos of 50-100 cell -stage and 100-130 minutes post-fertilization. 2 hours post-egg-laying, embryos were transferred using platinum wire onto glass slides with M9 solution. Embryos were washed 5 times using transfer-pipetting into M9 solutions, to remove bacteria before final mounting. These embryos of “late ball stage” (approximately 260-300 minutes post-fertilization) were mounted on a 2% agarose pad in H20 solution. 20-mm polystyrene beads as slide spacers in the sample preparation. This mounting causes stereotypical turns of the embryos in the eggshell during morphogenesis due to slight compression. Agarose pad was prepared using a 120 μm spacer on top of the slide to control the thickness. The final sample thickness was ~180 μm. Glass slides were covered with coverslips and sealed using liquid vaseline heated to 40°C, to prevent H20 evaporation. Brightfield and Brillouin images were acquired in 13-minute intervals.

#### Mouse organoids

Breeding and maintenance of the mouse colony was done in the Laboratory Animal Resources (LAR) facility of EMBL Heidelberg. Primary mammary epithelial cells were obtained from 8-week-old virgin females of FVB background. Three-dimensional (3D) cell cultures were established according to the published protocol^26,43^.

In short: The tissue from two mammary glands without mechanical dissociation was placed in 5-mL digestion medium (DMEM/F12 with L-glutamine, 15 mM HEPES (Biowhittaker, no. 12-719Q), supplemented with 1 M HEPES (Biowhittaker, no. 17-737E) to 25 mM final concentration, 150 U/mL Collagenase type 3 (Worthington, no. CLS3), and 20 μg/mLLiberase Blendzyme 2 (Roche, no. 11988425001), and digested for 15–16 h at 37°C in loosely capped 50-mL polypropylene conical tubes. The resultant organoid suspension was washed with 45 mL of phosphate-buffered saline (PBS) containing Ca^++^ and Mg^++^, pelleted at 1000 rpm for 5 min at room temperature, and resuspended in 5 mL of 0.25% trypsin-EDTA. After incubation for 40 min at 37°C in loosely capped 50-mL polypropylene conical tubes, cells were washed with 45 mL of DMEM/F12 with Lglutamine, 15 mM HEPES, supplemented with 1 M HEPES to 25 mM final concentration and with 10% Tet System Approved FBS (Clontech, no. 631101). Suspensions were treated with 10 mg/mL DNaseI (Sigma, no. D4527). Dissociated cells were pelleted at 1000 rpm for 5 min at room temperature and resuspended in PBS containing Ca^++^and Mg^++^, counted, and plated onto collagen-coated 10-cm plates (BioCoat, #356450) for overnight adhesion and expansion. Then cultured cells were washed with PBS without Ca^++^ and Mg^++^, and the remaining cells were treated with 1 mL 0.25% trypsin-EDTA. After cell detachment, trypsin was inactivated with 9 mL DMEM/F12 with Lglutamine, 25 mM HEPES, supplemented with 10% Tet System Approved FBS. Cells were pelleted at 1000 rpm for 5 min at room temperature and resuspended in PBS containing Ca^++^ and Mg^++^ and counted. Cells were mixed rapidly on ice with Matrighel Matrix (Corning, 354230) Droplets (100 μL) containing 10000 cells from mammary gland preparations and were dispensed in a petri dish (Mattek, 35mm Dish, No. 0 Coverslip) with a well in the center. After solidification on a level surface for 30 min at 37°C, the gels were placed at 37°C in a CO_2_ incubator with 1.5 mL of supplemented serum-free medium (500 ml MEBM mammary epithelial cell growth basal medium (Lonza, CC-3151) supplemented with 2 ml of bovine pituitary extract, 0.5 ml of hEGF, 0.5 ml of hydrocortisone, 0.5 ml of GA-1000, 0.5 ml insulin from MEGM mammary epithelial cell growth medium BulletKit (Lonza CC-3150)). Media was replaced every 2 days. A coverslip was used to cover the cell medium from evaporation and to allow access of the sample by two objectives.

#### Mouse strains and embryo culture

All mice used in this study were housed according to the guidelines of EMBL LaboratoryAnimal Resources. Mouse embryos were collected from superovulated (injection of 7.5international units (IU) pregnant mare serum followed by injection of 7.5 IU humanchorionic gonadotropin 42 to 50 hours later) 8- to 24-week-old female mice according to the guidelines of EMBL Laboratory Animal Resources. Embryos were cultured in 30 μl drops of G1 PLUS (Vitrolife) covered by mineral oil (Ovoil, Vitrolife). Embryos were isolated from C57BL/6J × C3H/He F1 females.

## Code availability

The GPU-accelerated SBS spectral analysis code will be made available at https://github.com/prevedel-lab/sbs-gpu-acceleration at the point of publication.

## Data availability

The raw datasets generated and/or analysed during the current study will be made publicly available at the point of publication.

