## Supplementary Information for "Pulsed stimulated Brillouin microscopy enables high-sensitivity mechanical imaging of live and fragile biological specimens"

### SUPPLEMENTARY NOTES

**SI Note 1: Practical limitations to achievable SNR enhancement through quasi-pulsing.**
Fig. 1 (e) and (f) show that 13 mW average pump power on the sample with 40 ns pulse width and 1.1 MHz repetition rate (duty cycle = 4.4%) has the same SNR as 295 mW CW pump. Therefore, we have demonstrated a pulse enhancement factor of 22.7 (i.e. 1/duty cycle). In our current setup, the narrowest pulse width is ~40 ns, which is limited by the AOMs. Further increases in the enhancement factor can be achieved by reducing the pulse repetition rate. However, this will also reduce the average power (and thus deteriorate the SNR) because of the limited peak pump and probe power. For a 40 ns pulsed setup, the maximum average probe power on the sample is limited to 5 mW because the input power to the fiber-coupled AOM of the probe beam is limited to ~650 mW to prevent damage. The probe AOM has 3 dB insertion loss and a 50:50 fiber coupler is used for coupling some of the probe light out as reference for the balanced detection. The maximum average pump power on the sample is limited to 13 mW for 40 ns pulsed setup. If one further reduces the pulse repetition rate, the average pump and probe power on the sample also decrease proportionally. This would affect the SNR, as the noise is dominated by the input electronic noise of the LIA.

To obtain a high shift and linewidth precision and hence a good Brillouin image, an SNR larger than 30 for water sample is typically needed. For example, as shown in Fig. 1(g), with 20 ms integration time, we use 60 ns pulse so that the probe power on the sample is 7 mW and on the photodiode is 5 mW. Compared to a 40 ns pulsed probe, the increase of the average probe power improves the SNR. The following parameters are used in all our Brillouin imaging experiments for the low power case: 60 ns pulse width, 1.1 MHz repetition rate, 20 mW average pump power and 7 mW average probe power impingent on the sample. This represents an enhancement factor of 15.2 and achieves the same SNR with 300 mW pump and 7 mW probe power for the CW case. It should be mentioned that in order to make a fair power comparison with a total power of ~265 mW[1], we use 700 ns pulse for pump with average power of 243 mW and for probe with average power of 7 mW. This setting is used in this work as 250 mW total power for a high-power case.

In our work, we used AOMs to generate pulses from CW lasers. This results in two limitations. Firstly, the pulse width is limited to several tens of nanoseconds. To generate shorter pulse width such as 1 ns, a fast electro-optic modulator (EOM) could be used. Secondly, the peak power and average power of the pulse is limited by the input power (i.e. the overall laser power). In the future, this could be solved by using an EOM to generate laser pulses at 1560 nm which could then be amplified by an Erbium-doped fiber amplifier (EDFA). The amplified pulse then propagates into a second-harmonic crystal (e.g. a PPLN waveguide) for second-harmonic generation so that a high power, short width 780 nm pulse is generated. A total enhancement factor of 10,000 could be achieved in this way by using 1 ns pulse train with 100 kHz repetition rate for the pump and probe, and 30 kHz (still in the low noise frequency range of the detection system as shown in SI Fig. 2) envelope pump modulation. This would allow it to obtain a similar SNR as the state-of-the-art CW scheme, but for 1 mW pump power and 20 us pixel time only.

**SI Note 2: SBS modulation frequency selection and optimal noise bandwidth/scan times.** SI Fig. 2 shows the noise spectra of a pulsed probe beam (90 ns pulse width, 1.1 MHz repetition rate) with and without balanced detection (i.e. differential input) measured by a lock-in-amplifier (MFLI, Zurich Instruments). It clearly shows that balanced detection can drastically decrease the noise. For example, when the probe average power on the photodiode is 12.3 mW, the noise density with balanced detection is 5.7 nV/sqrt(Hz) which is ~10 times smaller than with unbalanced detection at 320 kHz. The noise density is 2.5 nV/sqrt(Hz) when there is no probe light on the photodiode which is dominated by the input electrical noise of the LIA. With balanced detection, when the probe power on the photodetector is less than 6 mW, the low noise frequency band is flat <500 kHz. To minimize the input noise of LIA, the input signal must be filtered out at the frequency of the pulse repetition rate. Furthermore, the probe SBG gain at the frequency of the amplitude modulation of the pump needs to have low transmission loss. The pump envelope modulation frequency and the repetition rate of the pulse are selected to 320 kHz and 1.1 MHz so that a low pass filter (LPF-B0R35+, MiniCircuits) meets the above conditions.

The LIA is set to 2nd-order filter with a noise-equivalent bandwidth of 200 Hz with a frequency scan range of 2 GHz and a pixel time of 20 ms. This setting is used for all the cells, organoids, *C. elegans* embryo and mouse embryo imaging. For zebrafish larva imaging, the LIA is set to 2nd-order filter with noise-equivalent bandwidth of 100 Hz, a frequency scan range of 3 GHz and a pixel time of 40 ms. For *C. elegans* imaging in the head and gonad regions, the LIA is set to 2nd-order filter with noise-equivalent bandwidth of 300 Hz, a frequency scan range of 3 GHz and a pixel time of 20 ms. While our chosen frequency scan range of 2-3 GHz is slightly less than that of previous work[1], our simulations (SI Fig. 8) show that the spectral precision with 2-3 GHz scanning range is sufficiently high for both for single Lorentzian and double Lorentzian fitting.

In all the Brillouin imaging experiments, the pulsed-pump envelope modulation frequency is selected at 320 kHz. The pulsed probe beam is with 60 ns pulse width, 1.1 MHz repetition rate and 7 mW average power on the sample (corresponding to ~5 mW average power on the photodiode). The signal and reference beams are connected to the two differential inputs of the LIA with 50 Ω input impedance. For 200 Hz noise-equivalent bandwidth, the noise defined by the standard deviation of the LIA output when the pump-probe frequency difference is tuned away from the Brillouin peak, is measured to be 46.7 nV. Therefore, the measured noise density is 3.30 nV/sqrt(Hz). The average probe power on the photodiode is 5 mW and the responsivity of the photodiode is 0.55 A/W. So the shot-noise density of the single signal beam is $\sqrt{2e\eta P} * R = 1.5$ (nV/sqrt(Hz)), where $e$ is the elementary charge, $\eta$ is the responsivity of the photodiode (0.55 A/W for the photodiode we use), $P$ is the average power on the photodiode, $R$ is the input impedance of the LIA. The total estimated noise density, including the shot noise of the signal and the reference as well as the input electrical noise of the LIA, is calculated to be 3.28 nV/sqrt(Hz) which is very close to the measured noise. Note that the total noise is ~2 times larger than the shot noise of a single signal input.

**SUPPLEMENTARY FIGURES**

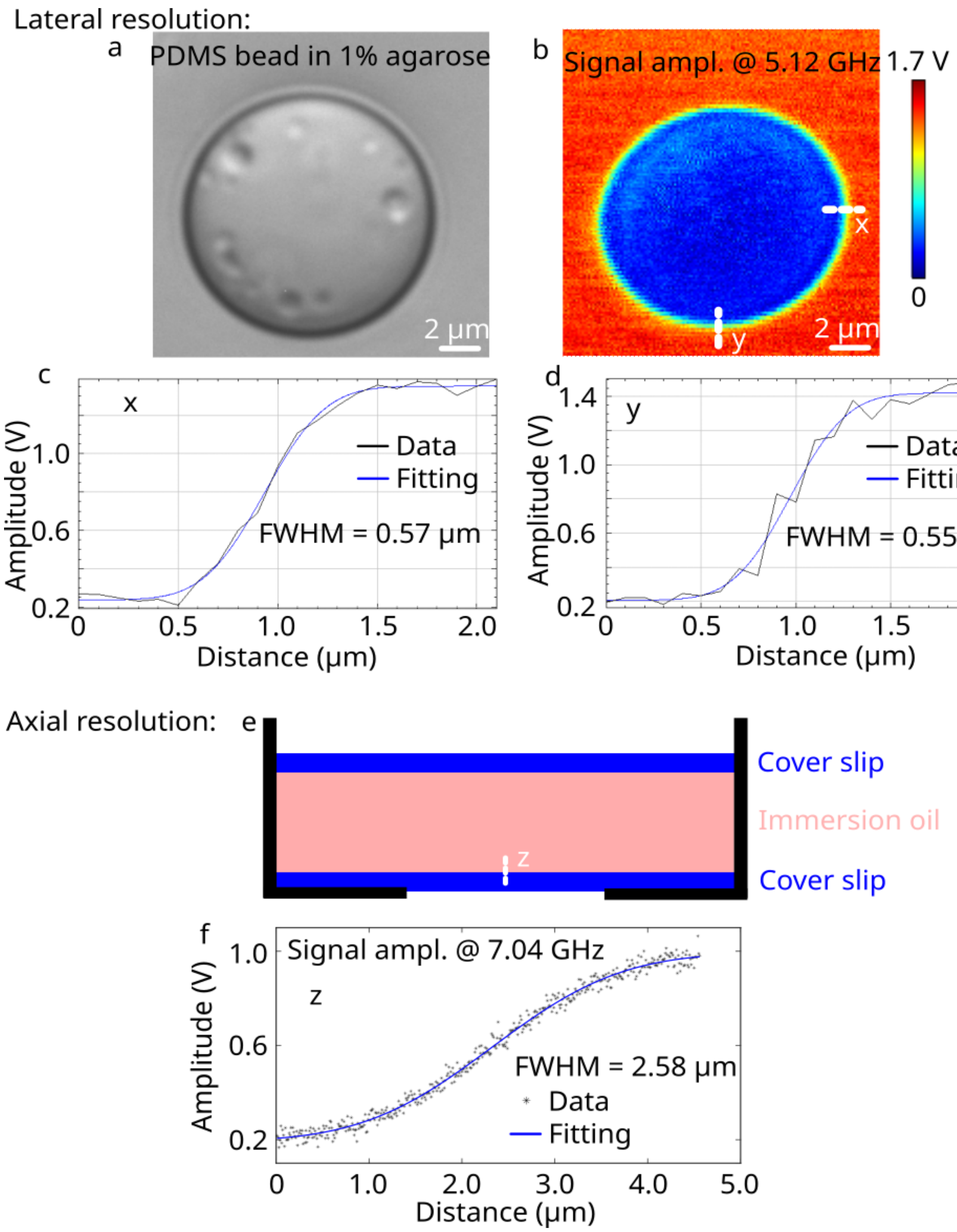


**SI Figure 1: Characterization of the lateral (x and y) and axial (z) resolution of the pulsed-SBS microscope.** The lateral resolution (x and y) and axial resolution (z) were measured with a PDMS bead in 1% agarose and immersion oil sandwiched by two cover slips, respectively. (**a**) Brightfield image of a PDMS bead in 1% agarose. (**b**) Fitted signal amplitude at 5.12GHz which is the Brillouin shift of 1% agarose. (**c**) and (**d**) are the amplitudes of Lorentzian fits to the spectrum, which are themselves fitted with an erf function at 5.12GHz along the edge in the x and y direction, respectively. (**e**) Schematic diagram of immersion oil sandwiched between two coverslips, with the white line indicating the measurement scan. (**f**) The amplitude of Lorentzian fits to the spectrum at 7.04GHz, which is the Brillouin shift of the immersion oil, and an erf-function fit to the data.

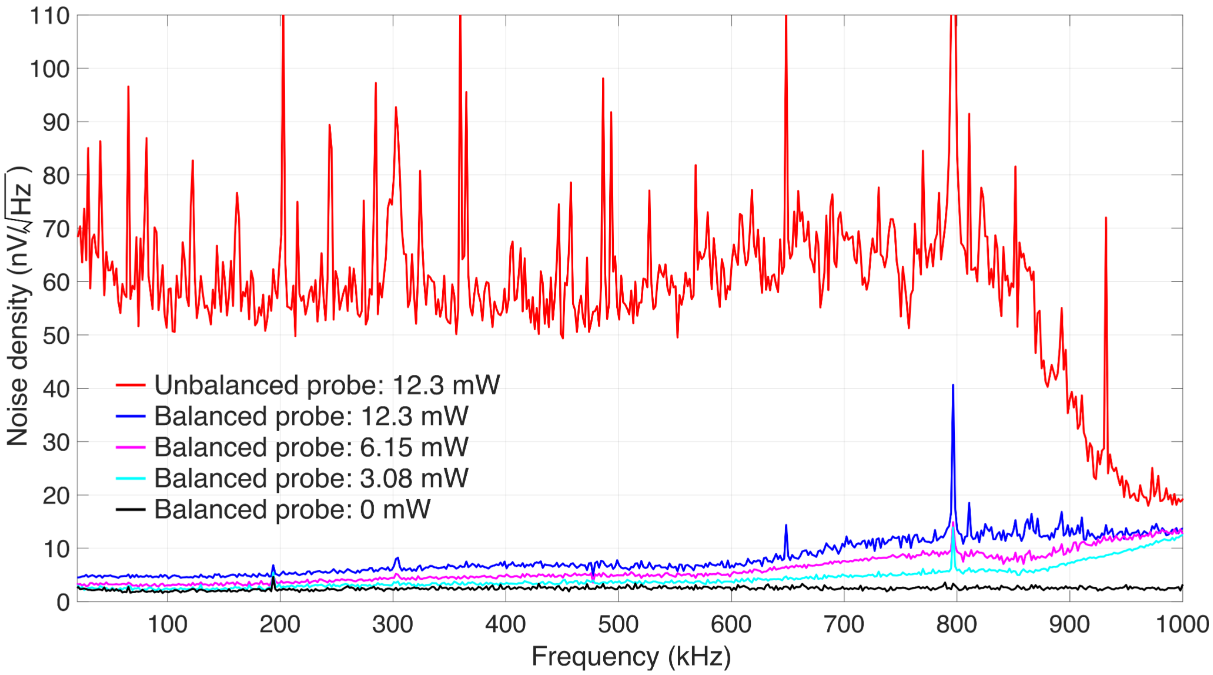


**SI Figure 2: Balanced vs. unbalanced detection performance of pulsed-SBS.** The plot shows the noise density against the frequency for different incident powers on the photodiode. Note that the pump amplitude envelope modulation frequency is selected at 320 kHz in our experiments.

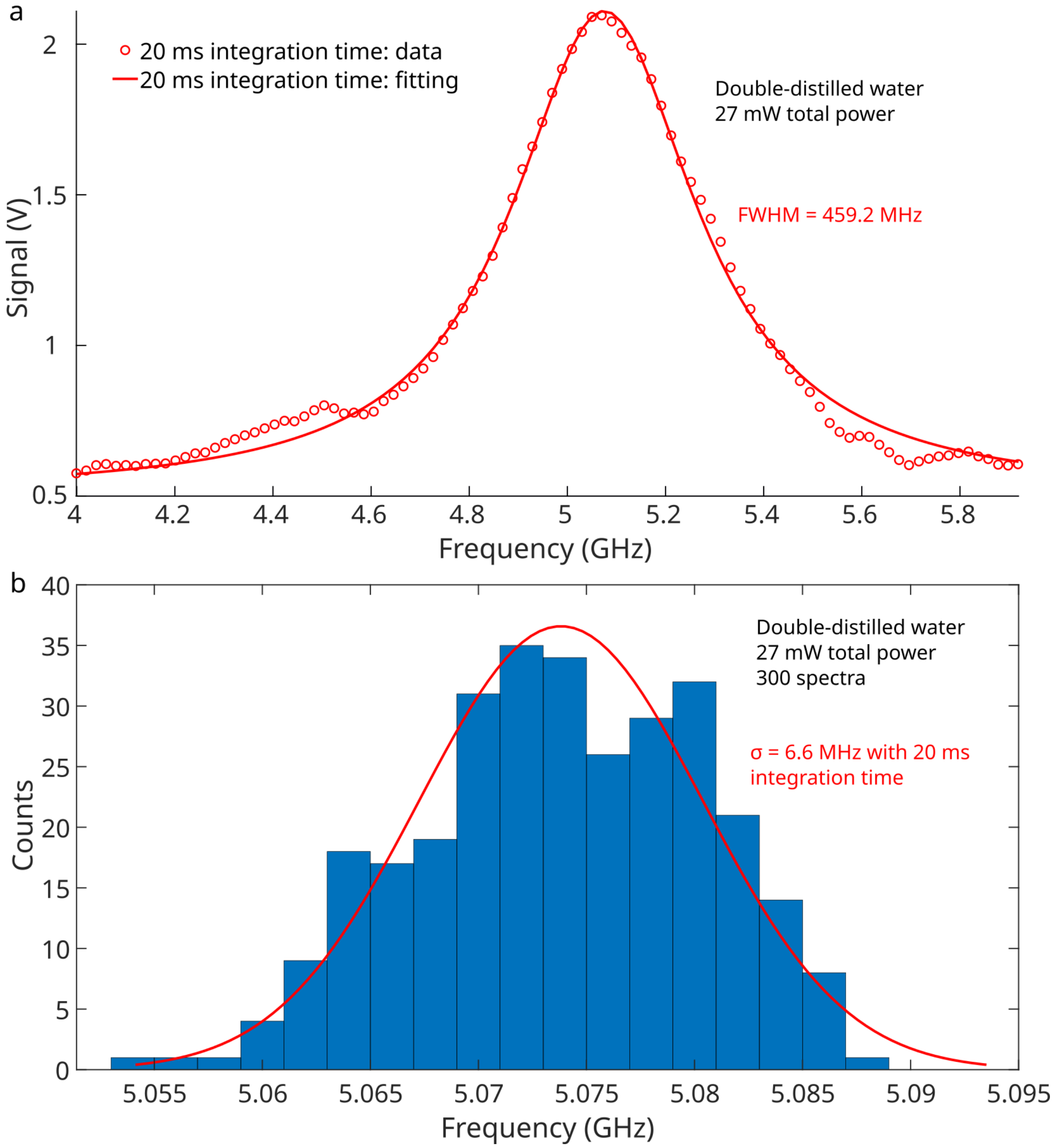


**SI Figure 3. SBG spectrum and measurement precision of water. (a)** A representative SBG spectrum of water with 20 ms integration time and 27 mW total power. **(b)** Brillouin shift precision, obtained from 300 sequential measurements of water with 20 ms integration time and 27 mW total power. These constitute the same parameters as used for most biological samples imaged in this work.

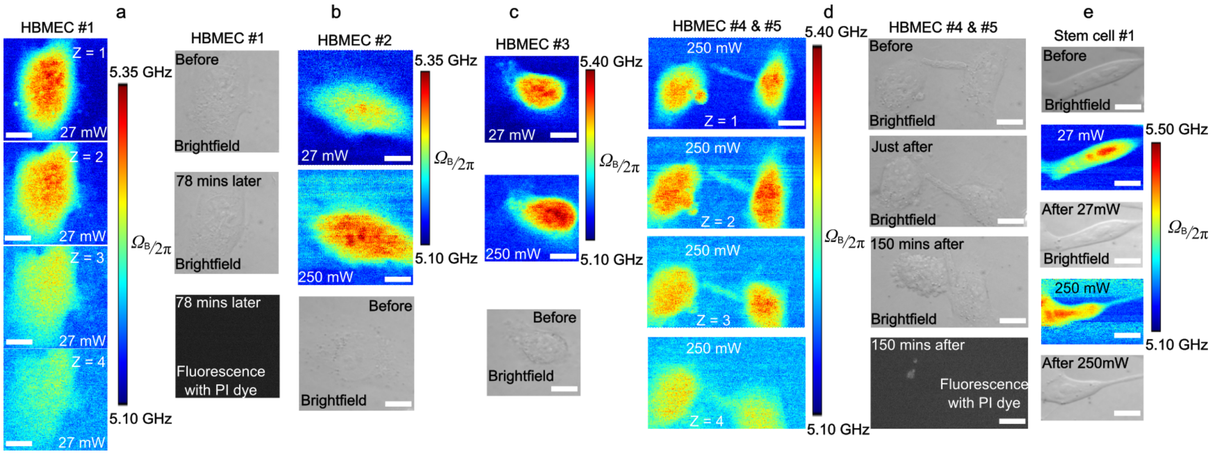


**SI Figure 4: SBS imaging of human brain microvascular endothelial cells (HBMEC) and a mouse embryonic stem cell. (a)** Three-dimensional sections of Brillouin shift images of a HBMEC under 20 mW pump and 7 mW probe with a z step of 1 µm. The co-registered brightfield image and fluorescence image of the cell with propidium iodide marker revealed no membrane damage 78 minutes after Brillouin imaging with 27 mW pulsed scheme. Brillouin shift images of HBMEC #2 (**b**) and HBMEC #3 (**c**) under 27 mW total power and 250 mW total power and the co-registered brightfield image. The overall Brillouin shift of the two cells were elevated due to high power as quantified in **Fig. 2j**. (**d**) Three-dimensional sections of Brillouin shift images of HBMEC #4 and #5 under 250 mW total power with a z step of 1 um. It showed direct effect of potential photodamage: First, clear cell blebbing (brightfield image) and membrane damage (fluorescence image) were observed in the CW scheme under 250 mW total power. Secondly, the right HBMEC showed increased motility which is indicative of cellular avoidance behavior. (**e**) A mouse stem cell was sequentially imaged with 27 mW pulsed and 250 mW CW schemes. The CW scheme resulted in motility of the stem cell. Furthermore, particle trapping and hence higher Brillouin shift in the cell medium is more evident in the CW scheme compared to pulsed-SBS. Scale bar: 10 µm.

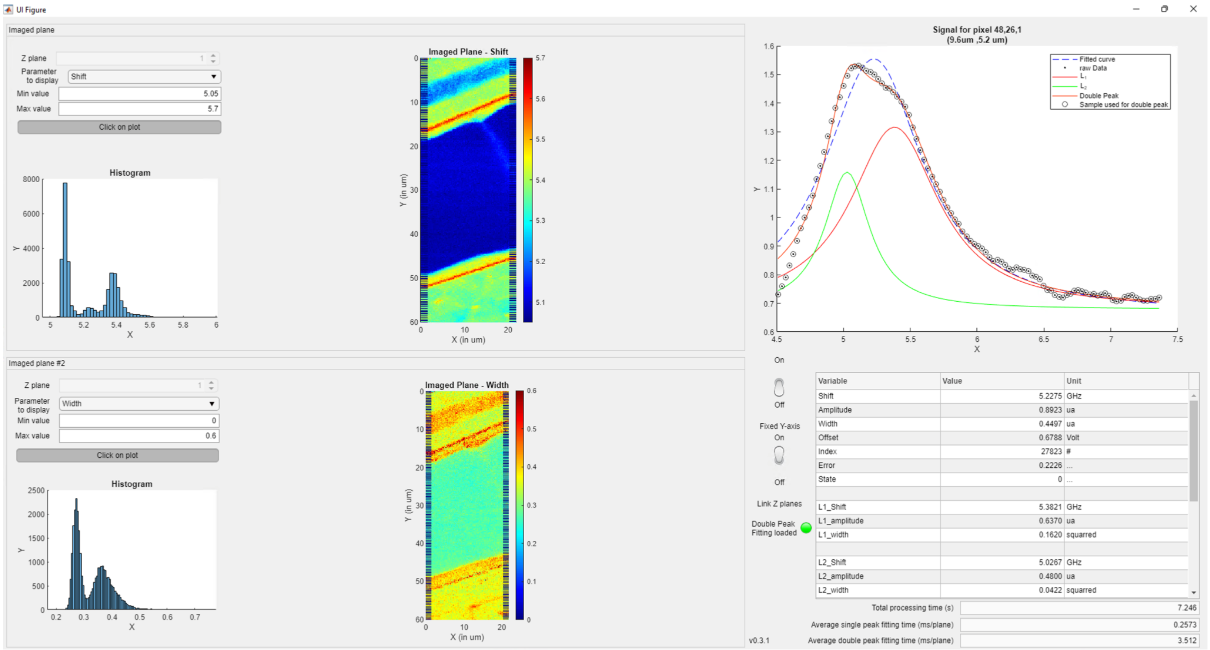


**SI Figure 5: Graphic user interface (GUI) of SBS spectral analysis.** The home-made GUI includes the parameter selection panel, 2 image panels after GPU fitting, histogram of the fitted parameter in the whole image, the raw data spectrum, the single peak fitting spectrum and the double peak fitting spectra as well as the fitted parameters at a specific pixel selected by the user. As an example, the pixel in the blood vessel at x = 9.6 µm, y= 5.2 µm position was selected and its SBG spectrum and the double-peak fitting results are shown in the GUI. Also see **Supplementary Software**.

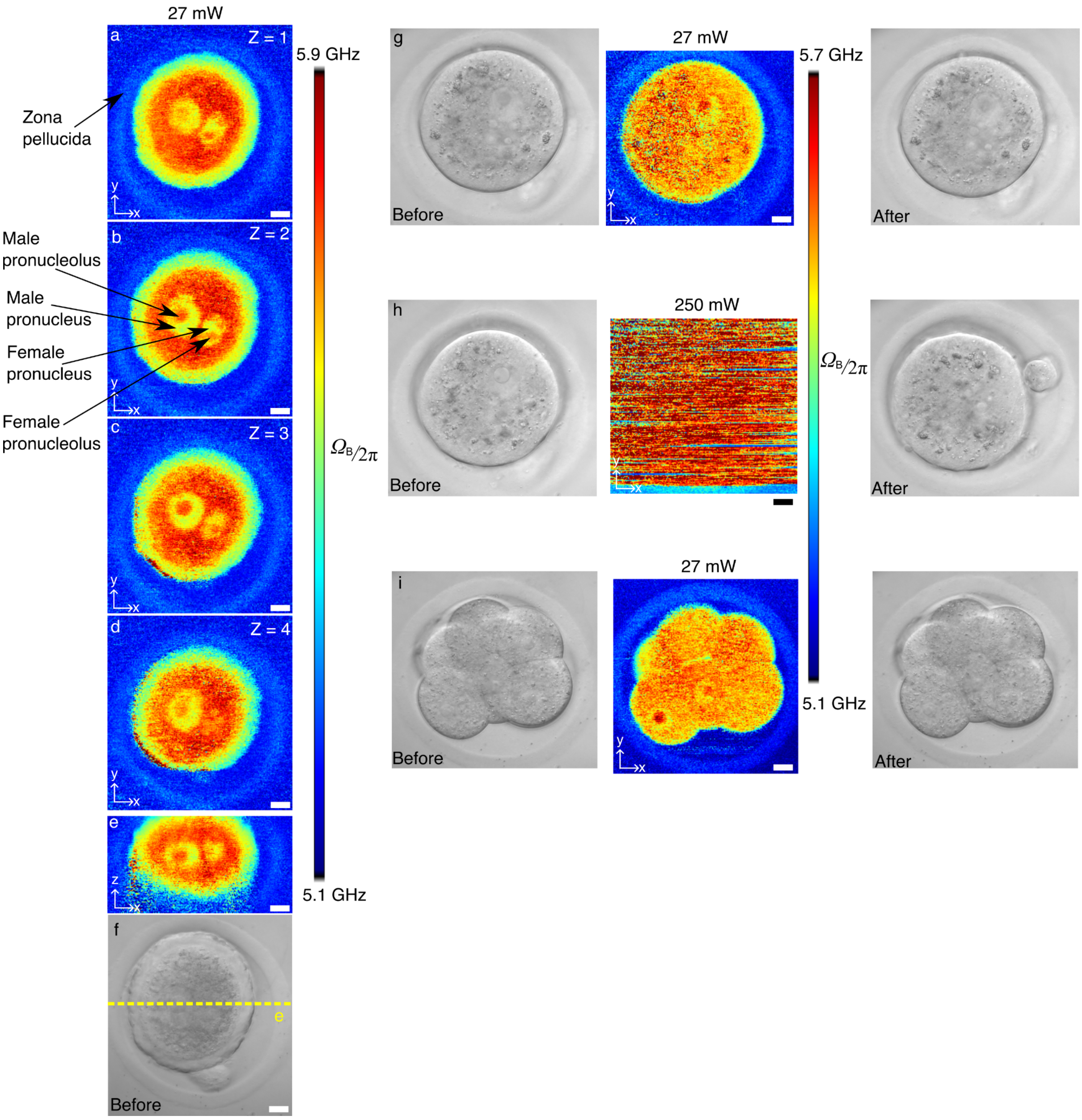


**SI Figure 6 | SBS imaging of mouse embryos.** Three-dimensional x-y cross sections(**a**)-(**d**) with a z step of 3 μm and (**e**) x-z cross sectional Brillouin images of a mouse zygote. (**f**) Brightfield image of the zygote. (**g**) Brillouin and brightfield images of a zygote under a 27 mW pulsed scheme. (**h**) Brillouin and brightfield images of a zygote under a 250 mW a CW scheme. Note that the high CW laser power trapped the zygote in the focus during the 2D piezo scanning. Therefore, it was not possible to obtain a Brillouin map of the embryo with the CW scheme. (**i**) Brillouin and brightfield images of an 8-cell-stage mouse embryo. Scale bar: 10 μm.

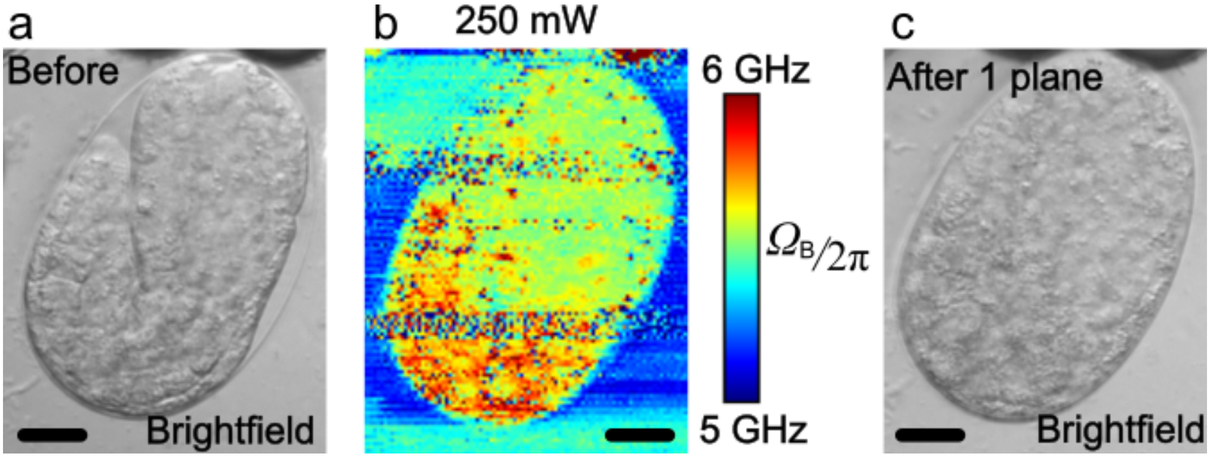


**SI Figure 7: Photodamage after CW-SBS imaging of *C. elegans* embryo. (a)** and **(c)** are the brightfield images of a *C. elegans* embryo before and after Brillouin imaging. (**b**) is the Brillouin image with 250 mW power in the CW-SBS scheme. Clear embryo death was observed in (**c**) after acquisition of a single Brillouin plane. The characteristic embryo morphology and individual cells of the 2-fold stage embryo (a) are lost after the Brillouin image acquisition (c). Scale bar: 10 μm.

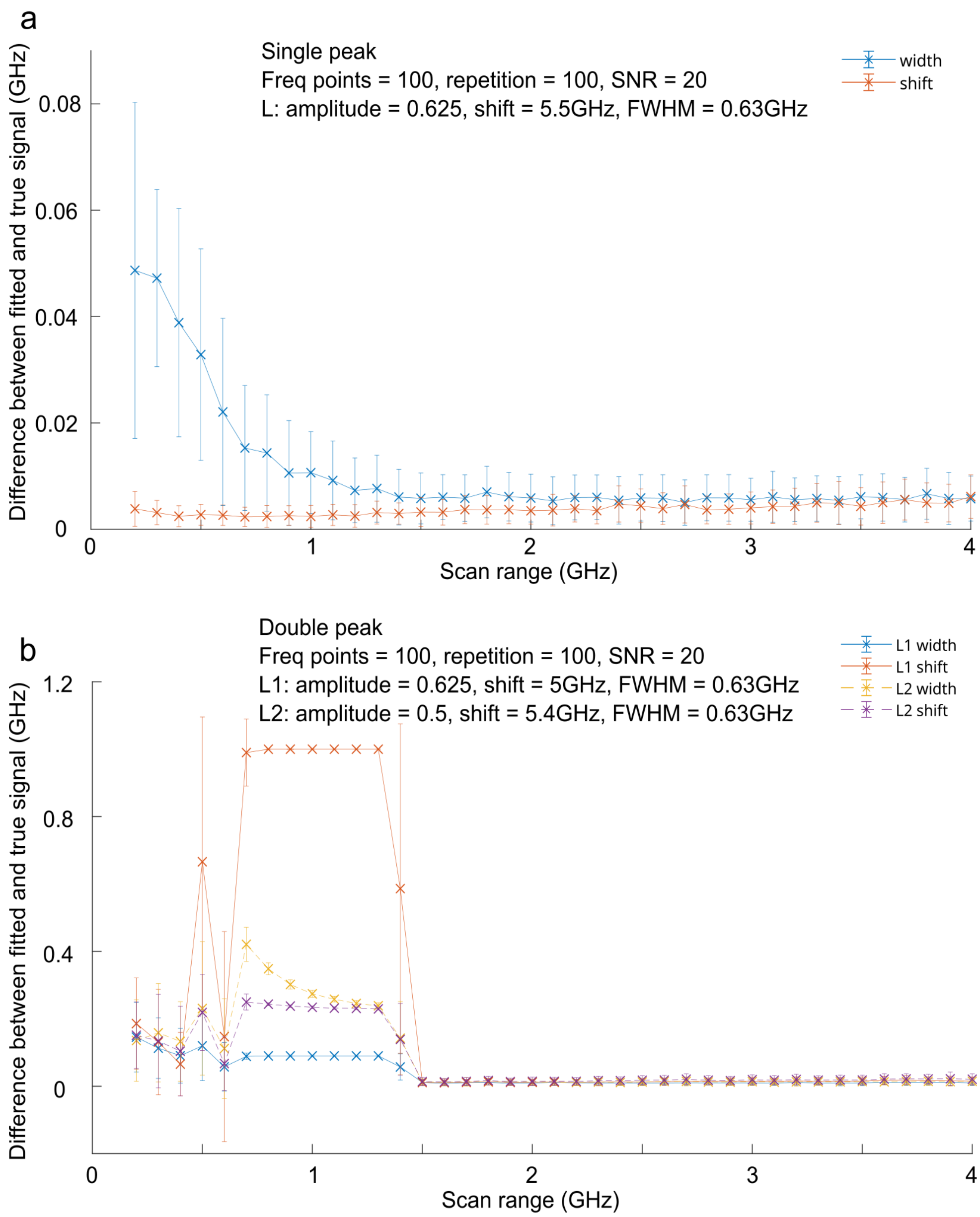


**SI Figure 8: Simulation of pulsed-SBS spectral precision under different scan ranges. (a)** The Brillouin shift and linewidth precision of single-peak fitting as a function of scan range. The signal to be fitted is the simulated sum of a Lorentzian (with an amplitude of 0.625, peak shift 5.5 GHz, FWHM 0.63 GHz) and a random noise with a standard deviation amplitude of 0.0312 (i.e. SNR of 20). **(b)** The Brillouin shift and linewidth precision of a double-peak fitting as a function of scan range. The signal to be fitted is the sum of two Lorentzian waveforms (L1 with an amplitude of 0.625, peak shift 5 GHz, FWHM 0.63 GHz, L2 with an amplitude of 0.5, peak shift 5.4 GHz, FWHM 0.63 GHz) and a random noise with a standard deviation of 0.0312 (i.e. SNR of 20). There are 100 frequency points in the scans, to mimic the actual experimentally acquired spectra. The precision and error bar are the mean and standard deviation of 100 independent simulations, respectively. Note that above 1.5 GHz, the double peak is fitted with L1 and L2, but between 0.8 and 1.5 GHz, only L2 gets a meaningful fit and L1 is minimized because it doesn't recognize the second peak.

| | CPU (Matlab) - Average of 7 repetitions | CPU + GPU (Gpufit) - Average of 10 repetitions | Speedup |
|---|---|---|---|
| Single peak: time per pixel/spectrum | 25.8 ms | 0.016 ms | x1541 |
| Double peak: time per pixel/spectrum | 68.1 ms | 0.088 ms | x775 |
| Total time to process 1 plane (30401 pixels total) | 2860.64s | 8.34 s | x343 |

**SI Table 1:** Processing times of the spectral dataset underlying Fig. 3c, which is composed of 301x101=30401pixels. The total processing time also includes additional steps, including loading of the data and computing fit statistics, which are currently only done on the CPU.

### SUPPLEMENTARY VIDEOS

**Supplementary Video 1: Time-lapse Brillouin shift maps of a developing *C. elegans* embryo.** Video corresponding to data shown in Fig. 5. Time is in minutes, scale-bar is 10µm.
